# Reproductive Trade-offs in Caribbean Corals: Plasticity Accelerates While Fragmentation Delays Reproductive Capacity

**DOI:** 10.64898/2026.05.14.725203

**Authors:** AC Vinton, C He, D Zdziebko, WC Million, R Cunning, E Bartels, EB Greenfield, CJ Krediet, CD Kenkel

## Abstract

In modular organisms, where growth and fragmentation blur the boundaries between individuals, the interplay between asexual and sexual reproduction creates complex fitness trade-offs. Life-history theory predicts that resources allocated to one fitness component necessarily reduce investment in others, yet detecting these trade-offs in wild populations of clonal organisms remains challenging. Phenotypic plasticity can enhance survival, yet its influence on reproductive capacity and life history trade-offs remains poorly understood. Using a fully crossed reciprocal transplant design, we tracked 263 colonies of the branching coral *Acropora cervicornis* across nine reef sites over 42 months, investigating relationships between fragmentation, morphological plasticity, and the capacity for sexual reproduction. Breakage patterns reflected both environmental and genetic factors. Primary branch breaks created a “double negative” effect—simultaneously more than doubling mortality risk and delaying attainment of a validated reproductive size class by ∼40%. Conversely, higher morphological plasticity in surface area-to-volume ratio accelerated sexual maturation up to 6-fold, counteracting the negative effects of fragmentation. In parallel, a simple demographic model parameterized with published fecundity data estimated that primary breakage reduces expected cumulative reproductive output by ∼58%, a result robust across a wide range of parameter assumptions. These results demonstrate a fundamental reproductive trade-off in which asexual reproduction through fragmentation undermines sexual reproductive potential by reducing colony size. Moreover, our findings reveal that fragmentation susceptibility is broadly heritable and subject to selection, and identify a compensatory mechanism through which plasticity enhances fitness beyond immediate survival.

## Introduction

Caribbean populations of the foundation coral *Acropora cervicornis* have declined by more than 90% since the 1970s (Lirman 2000), yet demographic data remain severely limited, with population assessments typically focused on coral cover rather than comprehensive tracking of individual colony outcomes over time (Goergen et al. 2019, 2020; Cant et al. 2022; Mercado-Molina 2015). This knowledge gap hinders the development of mechanistic models linking individual traits to population dynamics, particularly the trade-offs between sexual and asexual reproduction that may fundamentally determine population recovery trajectories.

Life history theory predicts fundamental trade-offs between growth, reproduction, and survival across diverse taxa (Stearns 1989; Roff 1992; Williams 1975). In modular organisms such as corals, where individual and population boundaries are blurred by growth and fragmentation, these trade-offs take on added complexity: asexual reproduction through fragmentation may reduce colony size below thresholds required for sexual reproduction, creating a conflict between the two reproductive modes (Williams 1975; Hughes et al. 1992; Okubo 2016).Phenotypic plasticity can modulate these trade-offs by enhancing fitness across heterogeneous environments (Scheiner 1993; DeWitt et al. 1998; Ghalambor et al. 2007), and was recently identified as adaptive in *A. cervicornis*, with more plastic individuals exhibiting superior growth and survival (Million et al. 2022). However, while plasticity can enhance immediate survival, its influence on reproductive capacity and the balance between sexual and asexual reproduction remains unknown (Vinton et al. 2022, 2023). This knowledge gap is critical because long-term reef recovery ultimately depends on successful sexual reproduction (Hughes & Tanner 2000). Understanding how plasticity shapes reproductive investment could reveal whether plastic responses primarily facilitate adaptation or create evolutionary constraints (Chevin et al. 2010; Chevin et al. 2013; Fox et al. 2019; Ghalambor et al. 2015; Vinton et al. 2022; Powell & Burgess 2025). Genetic variation in plasticity (GxE) that affects fitness was originally recognized as a key factor influencing reproductive mode in organisms with the capacity for clonal reproduction (Williams 1975), yet for reef-building corals specifically, we lack mechanistic understanding of how plasticity influences the balance between sexual and asexual reproduction—a relationship that fundamentally determines population recovery trajectories and adaptive potential in the face of increasing disturbance regimes.

Corals utilize both sexual and asexual reproductive strategies, although their relative contributions to population maintenance and recovery differ among species (Harrison & Wallace 1990; Highsmith 1982; Roff 2021). Sexual reproduction, particularly through broadcast spawning, offers two key advantages: greater dispersal distances essential for demographic rescue and novel genetic diversity crucial for adaptation to changing environments (Williams 1975; Drury & Lirman 2017; Harrison & Booth 2007; Chan et al. 2023). Asexual reproduction via fragmentation is thought to have been incorporated into coral life histories through natural selection, as indicated by morphological traits that promote, rather than resist, breakage (Highsmith 1982; Bak & Engel 1979). Genotype-environment interactions for fitness are prerequisites for adaptive evolution in organisms with mixed reproductive strategies, and maximizing clonal size is thought to directly enhance sexual reproductive capacity (Williams 1975). Although fragmentation was historically observed to contribute to rapid recovery following major disturbance events like storms (Shinn 1976; Gilmore and Hall 1976; Highsmith 1982); Caribbean reefs now demonstrate limited resilience (Gardner et al. 2003), likely due to chronically low sexual reproduction (Tunnicliffe 1981; Sammarco 1980; Dustan 1977; Tanner & Hughes 2000; Quinn 2005; Huntington et al 2011).

*Acropora cervicornis* exemplifies this dual reproductive strategy, undergoing annual mass spawning but also exhibiting high levels of asexual reproduction through fragmentation (breakage), a strategy that may represent an evolutionary adaptation to extreme weather events (Fong & Lirman 1995). The morphology and relatively fragile skeletal structure of these corals appear well-suited for storm-induced fragmentation, allowing broken pieces to establish new colonies following mechanical disturbance (Highsmith 1982; Roff 2021). This “storm-adapted” reproductive strategy has been harnessed by restoration programs that propagate and outplant coral fragments (Schopmeyer et al. 2017). Prior work has demonstrated differential asexual propagation among genetic lineages and within thickets of *A. cervicornis* (Tunnicliffe 1981; Drury et al. 2019). Nevertheless, the long-term fate and contribution of these fragments to sustaining populations remain largely unknown (Tunnicliffe 1981; Lirman 2000; Mercado-Molina et al. 2015). Tunnicliffe (1981) found that *A. cervicornis* fragments with multiple branches had higher survival rates than unbranched fragments, suggesting that morphological characteristics influence fragment viability.

Although early studies proposed that the rapid growth rates of branching coral species would favor both sexual and asexual reproduction with limited costs (Williams 1975; Highsmith 1982), more recent work suggests this dynamic could create a fundamental biological trade-off, as reproductive output in corals scales with colony size — larger colonies produce proportionally more gametes (Szmant 1986; Babcock 1991; Hall & Hughes 1996). Although post-pubescent corals may retain per-polyp fecundity even at reduced sizes (Rapuano et al. 2023), whole-colony reproductive output still depends on the total volume of gametogenic tissue. Coral mortality rates also exhibit strong size dependence, with substantial reductions once colonies surpass specific surface area thresholds—an observation that supports the hypothesis that fragmentation would only be adaptive in species with low mortality rates above certain size thresholds (Connell 1973; Highsmith 1982); while fragmentation, by reducing colony size, may delay or inhibit reproduction. Indeed, recent work has empirically confirmed that *A. cervicornis* must reach minimum size thresholds before achieving reproductive maturity, with Koch et al. (2022) directly validating reproductive competence in colonies exceeding ∼25 cm diameter (∼5,000 cm^3^ ellipsoid volume; equivalent to ∼161 cm total linear extension per the conversion in Boisvert et al. 2024 and Kiel et al. 2012). Although the occurrence and ecological significance of fragmentation have long been documented (Tunnicliffe 1981; Highsmith 1982; Shinn 1976), more recent experimental studies have treated breakage as unwanted noise, excluding fragmentation events from analyses of coral growth and survival—a methodological choice that overlooks a fundamental biological process likely critical to population dynamics (Highsmith 1982, Roff & Mumby 2012; Drury et al. 2019; Roff 2021). The combination of growth and fragmentation influences changes in whole colony morphology and the degree to which individual coral can change their morphology. In this study, we specifically quantify morphological plasticity through changes in surface area to volume ratio (SA:V), a single trait known to affect resource acquisition and colony growth rates which has been shown to vary among genotypes of *A. cervicornis* (Million et al. 2022, Conti-Jerpe et al. 2020). Beyond impacts on individual colony morphology, fragmentation generates important ecological consequences, including reef extension, patch reef formation, development of monospecific coral thickets, and the creation of habitat for diverse reef organisms (Highsmith 1982). Recent methodological advances now allow researchers to distinguish true growth from breakage (Million et al. 2022), facilitating quantification of the relative contributions of sexual and asexual reproduction and how they influence each other.

By employing time-to-event analyses on 263 fate-tracked coral colonies across multiple reef sites over 42 months we investigated three fundamental questions at the intersection of developmental plasticity and evolutionary biology: (1) What drives breakage patterns - are they random stochastic events or influenced by specific traits? (2) Does asexual reproduction through fragmentation create evolutionary trade-offs with sexual reproduction potential? and (3) How does morphological plasticity mediate these reproductive trade-offs? Our analyses reveal that breakage significantly extends time to reaching reproductive size in *A. cervicornis*, while morphological plasticity accelerates it. A demographic model parameterized with published fecundity data (Koch et al. 2022) estimates that primary breakage reduces expected cumulative reproductive output by ∼58%. Importantly, this reduction in sexual reproduction potential is not merely a site effect determined by environmental conditions; there is significant genotypic variation in susceptibility to breakage. Thus we verify the existence of a reproductive trade-off in *A. cervicornis* with profound implications for both evolution and conservation. Importantly, we demonstrate that breakage patterns are not merely random events driven by environmental stochasticity but are significantly influenced by coral genotype—a discovery that transforms breakage from experimental ‘noise’ into a broad-sense heritable trait subject to selection, challenging previous assertions that fragmentation results primarily from within-colony variation or environmental factors with no genetic basis (Highsmith 1982) and providing empirical evidence that fragmentation susceptibility itself may be subject to evolutionary processes.

## Methods

### Experimental Design and Field Monitoring

This study extends Million et al. (2022)’s 12-month dataset to 42 months to investigate putative reproductive milestones. The experimental design, field monitoring protocol, and 3D photogrammetry pipeline are summarized below; full details are provided in Million et al. (2022). Ten distinct *Acropora cervicornis* genotypes (designated as genotypes 1, 3, 7, 13, 31, 36, 41, 44, 50, and 62) were used in this study. These genotypes were originally collected from remnant populations in the lower Florida Keys between 2007-2012 and had been maintained through asexual propagation in common garden conditions at Mote Marine Laboratory’s in situ coral nursery at Looe Key for 5+ years prior to the start of this experiment. In April 2018, 270 coral ramets (27 per genotype, mean TLE 8.4 cm (TLE: total linear extension, defined as the sum of all branch lengths within a colony; Johnson et al. 2011)) were outplanted to nine active restoration sites across the Florida Keys reef tract at depths ranging from 5.6 to 9.1 m. These sites varied in environmental conditions, including wave exposure (with offshore sites experiencing greater oceanic swells), human visitation (ranging from minimal at remote sites to high dive tourism at sites with mooring buoys), and substrate characteristics. At each site, three ramets per genotype were randomly allocated to each of three ten-coral arrays. Ramets were attached to the reef substrate using a two-part marine epoxy (Z-Spar Splash Zone, A-788) applied to cleared areas of natural reef, following standard protocols for *A. cervicornis* outplanting (Johnson et al. 2011; Schopmeyer et al. 2017). No major bleaching events occurred during the study period (April 2018–October 2021). Ramets were photographed immediately before transplantation (following Million et al. 2022) and measured for TLE. They were resurveyed quarterly during the first year (July 2018, October 2018, January 2019, April 2019) then biannually for an additional 2.5 years (July 2019, October 2019, April 2020, October 2020, October 2021) to extend temporal scope and capture size-based indicators of reproductive development.

### 3D Photogrammetry and Trait Measurement

In situ photographs generated individual 3D models for each coral ramet using Metashape 1.5.4 (Agisoft LLC, St. Petersburg, Russia) following the high-throughput pipeline described in Million et al. (2021, 2022). Colony shape during the initial 12-month monitoring period was quantified using several metrics: SA:V (surface area-to-volume) and TLE-to-V ratios, packing, convexity, and sphericity (Million et al. 2022). This 12-month period captures the critical establishment phase when colonies undergo the most rapid structural development and morphological adjustment to site-specific environmental conditions (Million et al. 2022). We adopted the SA:V plasticity calculation from Million et al. (2022), who demonstrated that this morphological trait varies significantly among *A. cervicornis* genotypes, where plasticity is defined as the regression slope between genotype-specific SA:V values and site-specific population means, following the joint regression framework (Finlay and Wilkinson 1963; Pigliucci 2001; Murren et al. 2015). This unitless measure—values >1 indicate above-average plasticity—was calculated at four timepoints (T3, T6, T9, T12) through April 2019. After April 2019, morphology measurements ceased as corals became too large and structurally complex for reliable Meshlab quantification. Each genotype’s average SA:V plasticity value from the initial 12 months was used in subsequent survival and reproductive analyses. Breakage events were recorded at regular monitoring timepoints (T0, T3, T6, etc.) and classified by severity, branch location (primary, secondary, tertiary, quaternary), and damage extent (catastrophic or partial).

*A. cervicornis* reaches reproductive competence at colony sizes ≥25 cm diameter (∼5,000 cm^3^ ellipsoid volume), a threshold empirically validated by Koch et al. (2022) through direct observation of gamete bundles using replicate ramets of the same *A. cervicornis* genets used in this study. Using the TLE-to-ellipsoid volume conversion from Kiel et al. (2012; Equation 2 in Boisvert et al. 2024), this corresponds to approximately 161 cm TLE. Specifically, Koch et al. (2022) sampled branches from replicate ramets above this size threshold (≥25 cm diameter, equivalent to ∼5000 cm^3^ volume) from genets grown out in common garden nurseries (Looe Key and Sand Key) prior to the summer mass spawning window in years overlapping with our transplant study period (2020-2021). They visually confirmed the presence of gamete bundles in 80% of colonies in 2020 (n=61) and 100% in 2021 (n=39), thus validating this size-maturity relationship for the specific genotypes used in this analysis. We categorically assigned each coral as having reached a reproductive size class or not by manually reviewing 3D models (first 12 months) and representative 2D images (18-42 months) to determine whether colonies exceeded the 25-cm diameter threshold. Size was assessed using the square frame reference within each image (external dimensions: 24.76 cm × 24.76 cm). Coral whose branches extended beyond the square reference were recorded as exceeding the maturity threshold (1), while smaller colonies were recorded as immature (0, Fig. 1B).

**Figure 1.**
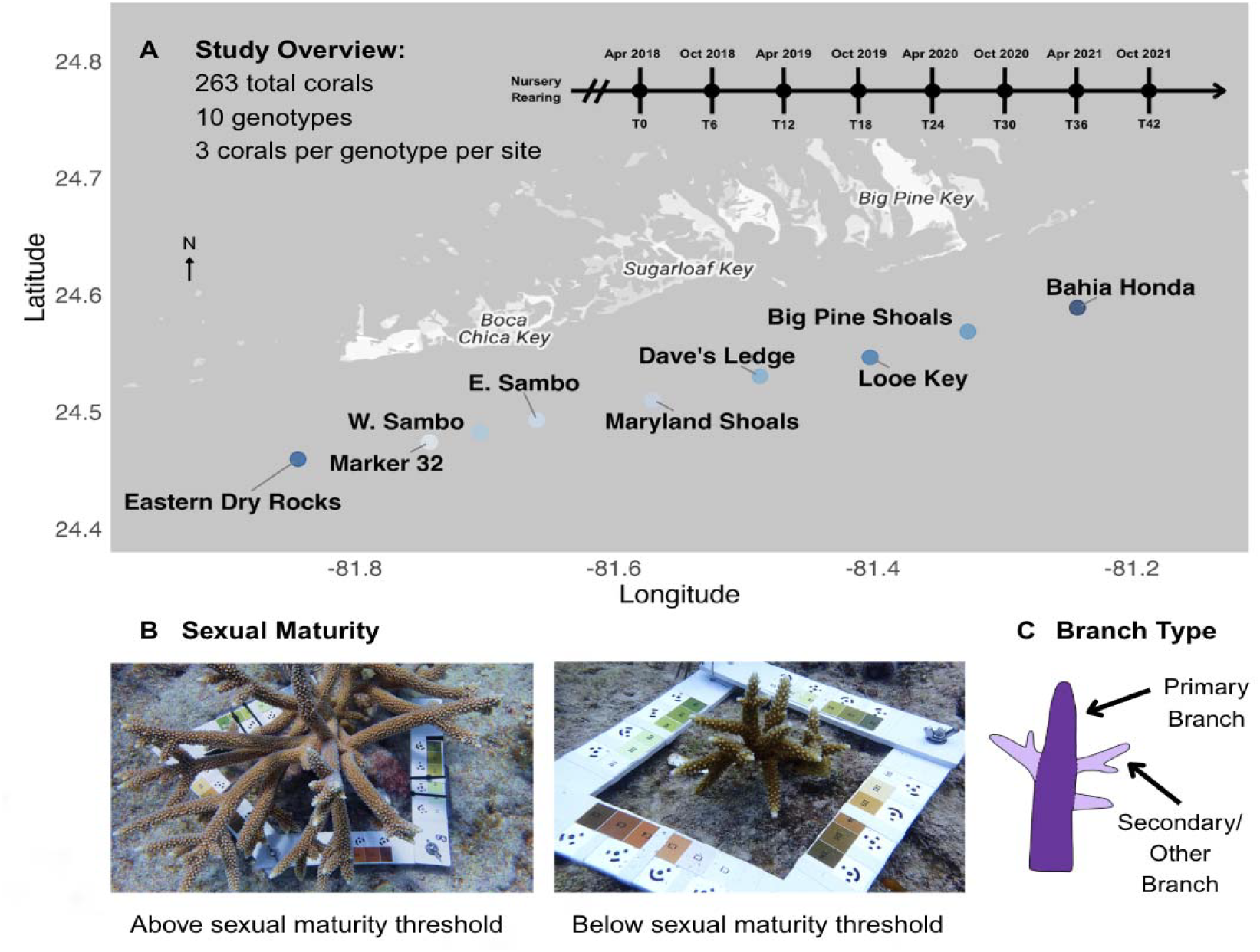
Study design and key morphological classifications for coral monitoring across the Florida Keys. (A) Study overview showing the spatial distribution of nine reef sites spanning the Florida Keys reef tract from Eastern Dry Rocks to Bahia Honda, with 263 total *Acropora cervicornis* colonies representing 10 genotypes (3 colonies outplanted per genotype per site). The timeline indicates monitoring timepoints from initial transplantation (T0) in April 2018 through 42 months (T42) in October 2021, with assessments conducted at 3-month intervals during the first year (T3, T6, T9, T12) and 6-month intervals thereafter. (B) Representative photographs showing coral colonies above and below the sexual maturity size threshold, demonstrating the visual differences in colony complexity and branch density associated with reproductive maturity. (C) Schematic illustration of branch hierarchy used for breakage classification, distinguishing between primary branches (main structural branches extending directly from the base) and secondary/other branches (smaller branches extending from primary branches).

### Statistical Analyses

All statistical analyses were performed in R (version 4.2.0), using the survival (Therneau 2023), survminer (Kassambara et al. 2021), coxme (Therneau 2022), and MuMIn (Bartoń 2023) packages. For detailed protocols regarding the experimental design and 3D photogrammetry pipeline, readers are referred to Million et al. (2022).

#### Variables and Definitions

Site: Nine reef locations where corals were outplanted (reference: E. Sambo)

Genotype: Ten distinct coral genotypes used in the study (reference: Genotype 36)

Initial colony size: Total linear extension (TLE; sum of all branch lengths) at the time of outplanting (T0)

Break types: Categorized by branch level affected (primary, secondary, tertiary, quaternary) and extent of damage (catastrophic or partial)

We analyzed several key variables to understand breakage patterns and their effects on survival and reproduction. Break type was classified based on which branch level was affected (primary, secondary, tertiary, or quaternary) and the extent of damage (catastrophic or partial). For simplified analyses, we created a binary classification distinguishing primary branch breaks from all other break types, as primary breaks had the most severe consequences for colony survival. Morphological plasticity was quantified as the genotype-level average of the joint regression slope of SA:V on site-mean SA:V, measured across the nine sites during the initial 12-month period (T3–T12; see Million et al. 2022 for derivation), with values above 1 indicating above-average plastic response to environmental conditions. We also tracked whether each colony experienced primary branch breakage at any point during the study, as this proved to be a critical predictor of both mortality risk and time to sexual maturity.

#### Analysis of Break Risk

To assess factors influencing coral breakage, we implemented recurrent event analysis using the Andersen-Gill extension of the Cox proportional hazards model. This allowed analysis of multiple breakage events per coral while accounting for within-colony correlation.

Predictors included genotype, site, and initial colony size. The coxph() function with Andersen-Gill extension treated each interval between monitoring points as a potential risk period. This realistically represented breakage processes as many colonies experienced multiple breaks.

Proportional hazards assumption was tested using Schoenfeld residuals. Performance was evaluated using concordance statistics, likelihood ratio tests, and Wald tests. We calculated hazard ratios with 95% confidence intervals for each covariate. Kaplan-Meier curves visualized breakage-free probability over time across genotypes and sites, with log-rank tests for statistical significance.

#### Survival Analysis

To analyze coral survival, we used Cox proportional hazards models with site, genotype, and simplified break type as predictors. Models were fitted using coxph() with reference categories defined above. We tested the proportional hazards assumption using Schoenfeld residuals and calculated hazard ratios with 95% confidence intervals for each covariate.

To test whether morphological plasticity modulates the effects of breakage on survival, we conducted stratified analyses examining the interaction between break type and plasticity level. Corals were classified as high or low plasticity based on median SA:V plasticity (median = 0.87), and the interaction between simplified break type and plasticity group was included in the Cox model.

#### Time to Sexual Maturity Analysis

Cox proportional hazards models examined how initial size, primary break presence, and SA:V plasticity influenced sexual maturity timing across reef sites.

To understand how multiple factors jointly influence the potential to achieve a size exceeding the sexual maturation threshold (≥ 25-cm diameter), we tested whether the effects of initial size, breakage, and plasticity varied across sites. We started with a comprehensive model including all possible interactions, then systematically removed non-significant terms to identify the most important relationships while maintaining model interpretability. Models were compared using AIC, likelihood ratio tests, Wald tests, and concordance statistics.

We calculated hazard ratios with 95% confidence intervals and tested proportional hazards assumptions using Schoenfeld residuals. Additionally, we compared time to maturity between high and low plasticity groups among corals experiencing primary breaks to test whether plasticity enhances post-breakage recovery.

##### Expected Reproductive Output Model

To estimate the demographic consequences of fragmentation for reproductive output independent of any binary maturity threshold, we constructed a simple demographic model following Koch et al. (2022). Expected cumulative reproductive output was calculated as:

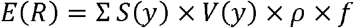

where *S*(*y*) is the Kaplan-Meier survival probability at annual spawn event *y, V*(*y*) is colony volume estimated from TLE using an empirical power-law fit to our photogrammetry data, ρ is polyp density (16 polyps/cm^2^), and *f* is oocytes per polyp (7 oocytes/polyp), with parameter values from Koch et al. (2022, Table 1). Colony growth was projected using von Bertalanffy curves (TLE(*t*) == *L∞* − (*L∞* − *L*_0_) X e^−kt^) fit to individual-level TLE data from T0–T12 separately for colonies with and without primary breaks, with *L∞* bounded at 200 cm (no primary break) and 150 cm (primary break) based on observed maximum TLE values and field observations (van Woesik et al. 2021; Sánchez et al. 2025). Expected reproductive output was summed across three annual spawn events (T12, T24, T36), corresponding to the August mass spawning of *A. cervicornis* in the Florida Keys. This approach assumes fecundity scales continuously with colony volume (Hall & Hughes 1996; Álvarez-Noriega et al. 2016), avoiding the need for a binary maturity threshold. We assessed model sensitivity by varying all parameters across their published ranges and tested robustness to the choice of L∞ by fitting models with asymptotes ranging from 100–300 cm TLE. The reproductive threshold from Koch et al. (2022; ∼5,000–10,000 cm^3^ ellipsoid volume, equivalent to ∼161–249 cm TLE via Boisvert et al. 2024) is shown for reference but does not enter the model calculation.

## Results

### Fragmentation Pattern Analysis

Analysis of 263 unique corals revealed that 83.7% (220 corals) experienced at least one breakage event during the study period. Breakage events were categorized by branch location (primary, secondary, tertiary, quaternary) and extent (catastrophic vs. partial). The most common fragmentation type was Primary Catastrophic (code “1C”), accounting for 67.3% of all breaks, while 16.0% of corals remained intact throughout the observation period.

Breakage patterns varied significantly among genotypes. Notably, Genotype 13 had the highest break rate with 100% of corals experiencing breaks, followed by Genotype 41 (92.3%) and Genotypes 3 and 7 (both approximately 89%). In contrast, Genotypes 50 and 36 showed the lowest breakage rates (73.1% and 74.1%, respectively). Site analysis showed high break rates at Bahia Honda (96.6%) and Dave’s Ledge (96.4%), while E. Sambo and Marker 32 had the lowest break rates (both 66.7%).

Kaplan-Meier survival analysis demonstrated significant differences in breakage-free probability over time among both genotypes (log-rank p < 0.001) and sites (log-rank p < 0.001) (Fig 2B, C). The probability of corals remaining intact declined most rapidly in the first 9 months, with only 37.1% of corals remaining unbroken by this time point, further decreasing to 20.2% by month 18.

**Figure 2.**
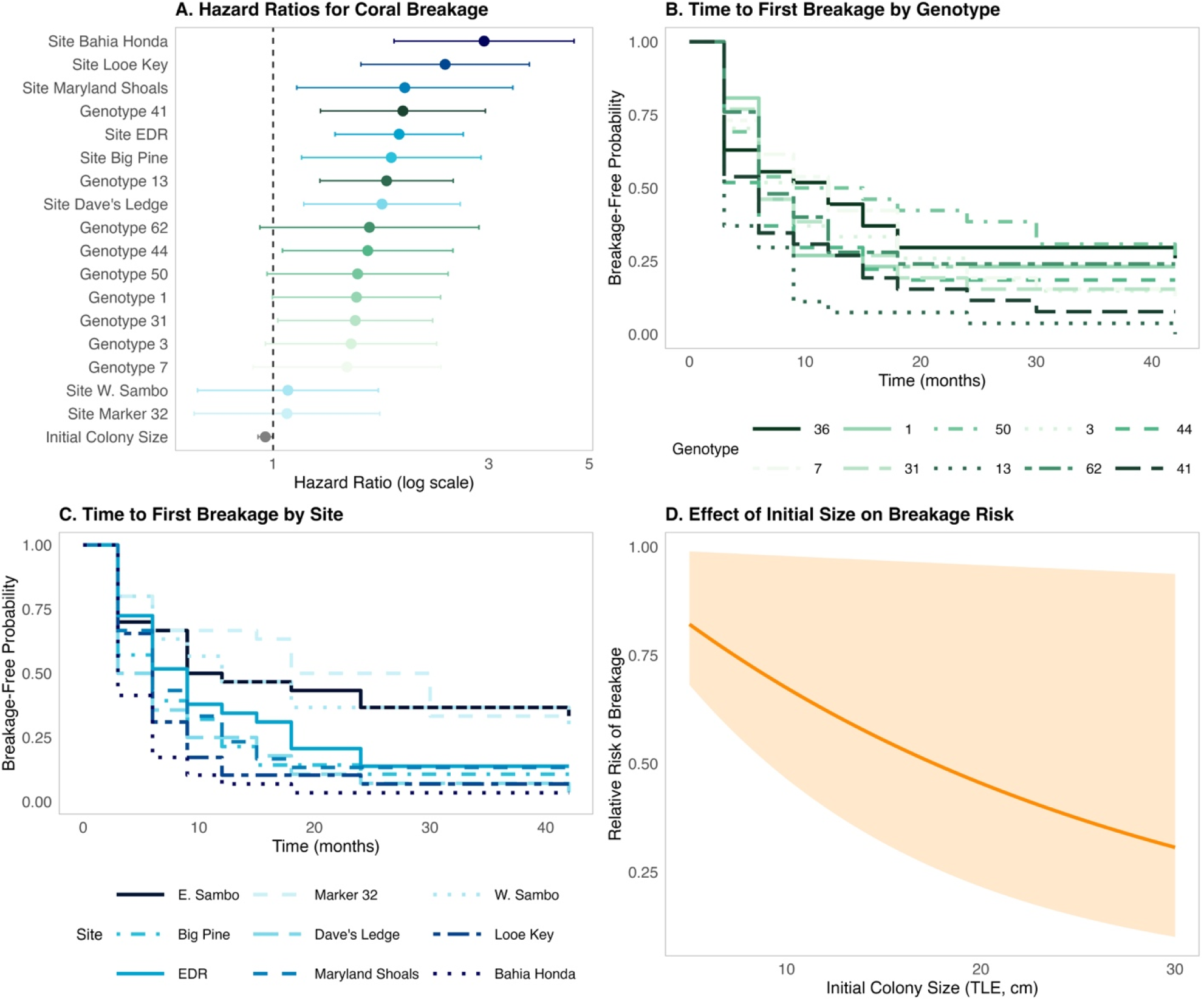
Multi-panel Analysis of Coral Breakage Risk Factors. Factors influencing coral breakage risk derived from recurrent event Cox proportional hazards modeling (Cox, 1972; Andersen & Gill, 1982). Panel A displays a forest plot of hazard ratios (HR) with 95% confidence intervals from an Andersen-Gill recurrent event Cox proportional hazards model examining factors associated with coral breakage events. The model incorporates genotype (reference: Genotype 36), site (reference: E. Sambo), and initial coral size as predictors, while accounting for within-subject correlation using clustering by coral tag. The vertical dashed line at HR=1 represents the reference point (no effect). Predictors with hazard ratios greater than 1 indicate increased risk of breakage events, while hazard ratios less than 1 suggest reduced risk. Variables are color-coded by type: genotypes in shades of green, sites in shades of blue, and initial size in orange, with color intensity corresponding to the magnitude of effect. Variables are ordered by magnitude of hazard ratio, with those having the strongest association with increased breakage risk at the top. Panel B presents Kaplan-Meier curves showing time to first breakage by genotype (Kaplan & Meier, 1958). Each colored line represents the survival function (breakage-free probability) for a different coral genotype over time (months). These curves illustrate the varying resistance to breakage among different genetic lineages throughout the monitoring period. Panel C displays Kaplan-Meier curves for time to first breakage by site location. The distinct colored lines represent different reef sites, with E. Sambo as the reference site, showing how environmental conditions at each location influence breakage patterns over time. Sites with curves that remain higher have lower breakage rates, while steeper declining curves indicate sites with higher breakage frequencies. Panel D illustrates the effect of initial colony size on breakage risk. The orange line shows the modeled relationship between a coral’s size at outplanting and its relative risk of experiencing a break, with the shaded region representing the 95% confidence interval.

### Recurrent Break Event Analysis

The Andersen-Gill Cox model for recurrent break events showed a concordance of 0.625 (se = 0.015) and satisfied the proportional hazards assumption (global p = 0.42). Genotype hazard ratios were significantly elevated for Genotype 13 (HR = 1.78, 95% CI: 1.27-2.50, p < 0.001), Genotype 41 (HR = 1.94, 95% CI: 1.27-2.95, p = 0.002), Genotype 44 (HR = 1.62, 95% CI: 1.05-2.50, p = 0.030), and Genotype 31 (HR = 1.52, 95% CI: 1.02-2.25, p = 0.038) relative to the reference genotype 36 (Fig 2A).

Site location significantly influenced coral breakage risk. The Andersen-Gill Cox model for recurrent break events (concordance = 0.625, se = 0.015) revealed substantial variation in breakage hazard across sites relative to E. Sambo, the reference site, which experienced 66.7% breakage. E. Sambo was selected as the reference site because it had the lowest breakage rate, providing a conservative baseline for comparison; results were qualitatively robust to alternative reference site choices. Sites with the highest breakage risk were Bahia Honda (HR = 2.93, 95% CI: 1.85-4.63, p < 0.001) and Looe Key (HR = 2.40, 95% CI: 1.56-3.68, p < 0.001), where corals were 2.4 to 2.9 times more likely to break than at E. Sambo. Four additional sites (Maryland Shoals, EDR, Big Pine, and Dave’s Ledge) showed significant increases in breakage risk (HR range: 1.74-1.95, all p < 0.02), while Marker 32 and W. Sambo showed no significant difference from E. Sambo (Fig 2A). Initial size showed a small but significant protective effect against subsequent recurrent breaks (HR = 0.96, 95% CI: 0.93-1.00, p = 0.038) (Fig 2D).

### Survival Analysis

Our Cox proportional hazards model examining the effects of site, genotype and break type on coral survival demonstrated that primary branch breaks significantly increased mortality risk (HR = 2.58, 95% CI: 1.14-5.82, p = 0.022), while secondary or other breaks showed a potential protective effect, though it was not statistically significant (HR = 0.20, 95% CI: 0.02-1.70, p = 0.142) (Fig 3A, D).

**Figure 3.**
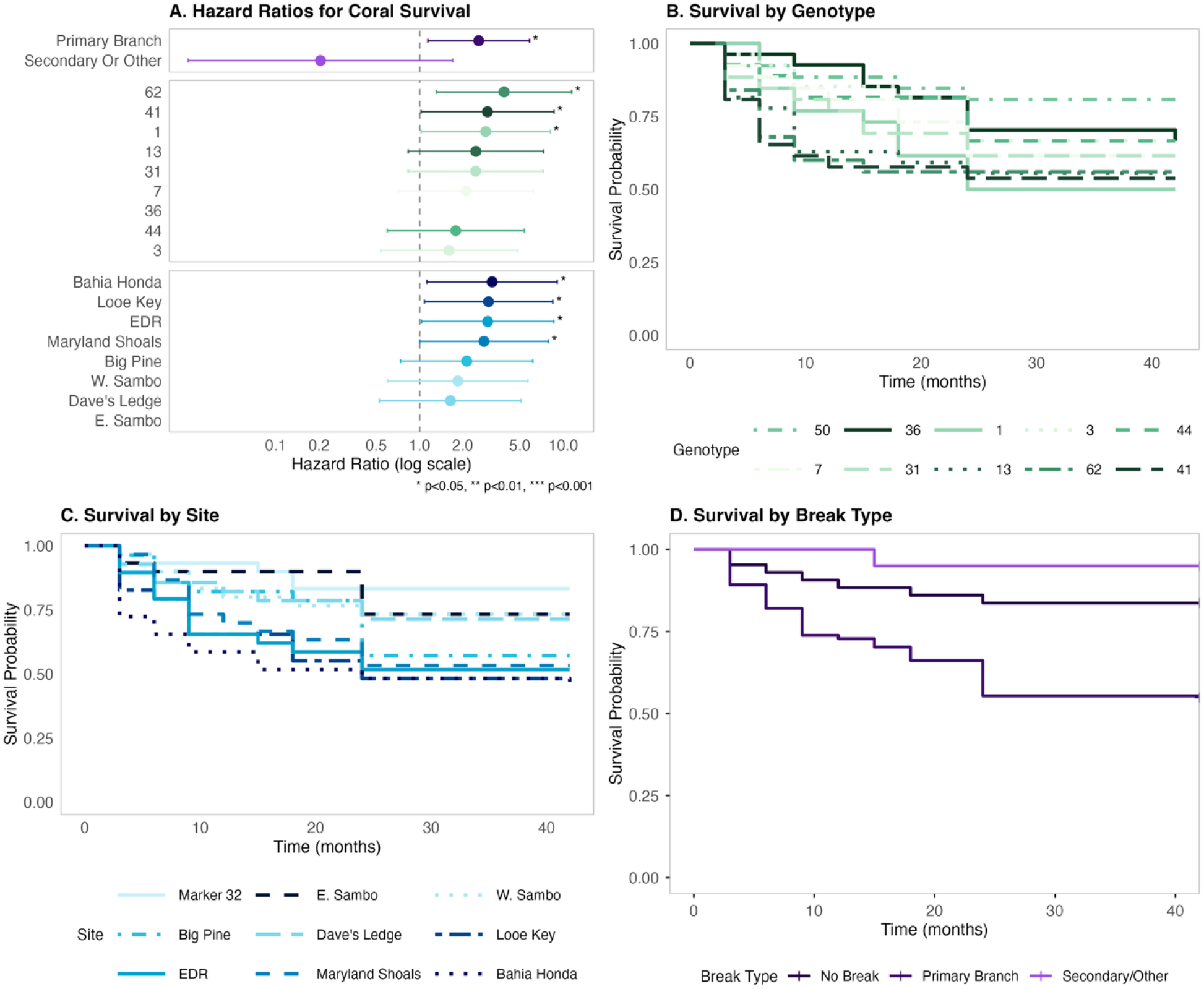
Factors affecting coral survival. **Panel A** shows a forest plot of hazard ratios from a Cox proportional hazards model incorporating site (reference = Marker 32), genotype (reference = 50), and break type (reference = No Break) as predictors. Hazard ratios greater than 1 (to the right of the dashed line) indicate increased mortality risk, while values less than 1 indicate reduced risk. Variables are color-coded by category (genotypes in green, sites in blue, break types in purple), grouped by category, and ordered by effect size within each group, with statistical significance indicated by asterisks. **Panel B** displays Kaplan-Meier survival curves stratified by genotype, with each genotype shown in a different shade of green. **Panel C** shows survival curves stratified by site, with sites displayed in shades of blue, revealing significant site-specific differences in survival probability over time. **Panel D** presents survival curves by break type (No Break, Primary Branch, Secondary/Other) in shades of purple, highlighting how different break types influence subsequent survival outcomes.

Site effects on mortality risk were significant for Looe Key (HR = 3.02, 95% CI: 1.08-8.45, p = 0.035), EDR (HR = 2.98, 95% CI: 1.03-8.60, p = 0.043), Maryland Shoals (HR = 2.81, 95% CI: 1.00-7.88, p = 0.050), and Bahia Honda (HR = 3.20, 95% CI: 1.13-9.07, p = 0.029) (Fig 3A, C). Genotype effects were significant for Genotypes 62 (HR = 3.87, 95% CI: 1.31-11.44, p = 0.014), 41 (HR = 2.97, 95% CI: 1.02-8.63, p = 0.045), and 1 (HR = 2.89, 95% CI: 1.03-8.13, p = 0.045) (Fig 3A, B).

The model showed good concordance (0.692, SE = 0.026) and satisfied the proportional hazards assumption (global test: p = 0.104), indicating valid statistical inference.

To test whether morphological plasticity modulates the effects of breakage on survival, we conducted a stratified analysis examining the interaction between break type and plasticity level. Corals were classified as high or low plasticity based on median SA:V plasticity (median = 0.87). The interaction model revealed that secondary breaks had markedly different effects depending on plasticity level. In high-plasticity corals, secondary breaks were strongly protective (HR = 1.25 × 10^-^□, p = 0.996), while in low-plasticity corals the protective effect was weaker (HR = 0.29). Notably, high-plasticity corals with secondary breaks experienced 0% mortality (0/7) compared to 10.5% mortality (2/19) in high-plasticity corals without breaks. Low-plasticity corals showed 7.7% mortality (1/13) with secondary breaks versus 20.8% (5/24) without breaks (Table S9). In contrast, primary breaks increased mortality risk regardless of plasticity level, though the effect was stronger in high-plasticity corals (HR = 3.83, 95% CI: 0.90–16.28, p = 0.069) than low-plasticity corals (HR = 2.13). The interaction between primary breaks and plasticity was not significant (p = 0.50).

### Time to Sexual Maturity Analysis

After excluding corals with missing breakage information or incomplete time-to-event records, a total of 177 colonies were included in the analysis. Colonies were classified as having experienced a primary break if their most severe breakage was recorded as a primary catastrophic or primary partial break. This filtered dataset ensured complete and consistent information on initial size, surface area to volume plasticity, breakage history, and time to sexual maturity. Of the 177 corals analyzed for sexual maturity, 76 (43%) reached a mature size class during the observation period. The Cox proportional hazards model with interactions demonstrated strong statistical significance (likelihood ratio test: χ^2^(20) = 72.68, p = 7e-08) and excellent concordance (0.813, SE = 0.026).

Initial size showed a significant positive effect on the hazard of maturing (HR = 1.390, 95% CI: 1.134-1.703, p = 0.002), indicating that larger colonies at T0 matured more quickly. Corals with primary breaks had significantly delayed maturation (HR = 0.570, 95% CI: 0.337-0.962, p = 0.035). Higher average plasticity in surface area to volume ratio in the first twelve months also significantly accelerated maturation (HR = 6.556, 95% CI: 1.979-21.714, p = 0.002). While the model included site-by-initial size interactions, none reached statistical significance (all p > 0.05), suggesting the beneficial effect of larger initial size on maturation was consistent across sites.

The model revealed a significant interaction between the initial size of the coral fragment and the coral’s plasticity in surface area to volume ratio (HR = 0.837, 95% CI: 0.738-0.951, p = 0.006), indicating that the accelerating effect of morphological plasticity on maturation was more pronounced in smaller initial fragments. Site-specific effects were evident, with Marker 32 (HR = 45.792, 95% CI: 4.383–478.433, p = 0.001) exhibiting significantly faster maturation rates compared to the reference site, E. Sambo, suggesting strong environmental influences on developmental timing at this location despite standardized genotypic composition.

Kaplan-Meier survival analyses revealed substantial variation in time to sexual maturity across sites, with corals at Marker 32 maturing significantly faster than those at other locations, particularly E. Sambo (Fig. 4B). Primary branch breakage significantly reduced the probability of reaching sexual maturity over time, with corals that experienced primary breaks remaining immature longer than intact individuals (Fig. 4D). Initial colony size also influenced maturation timing, with larger corals reaching maturity more quickly, as indicated by a steeper decline in the proportion of immature colonies over time compared to smaller size classes (Fig. 4C). Morphological plasticity showed a strong positive relationship with maturation rate, with highly plastic corals maturing up to 6x faster than those with low plasticity across the observed range (Fig. 4E).

**Figure 4.**
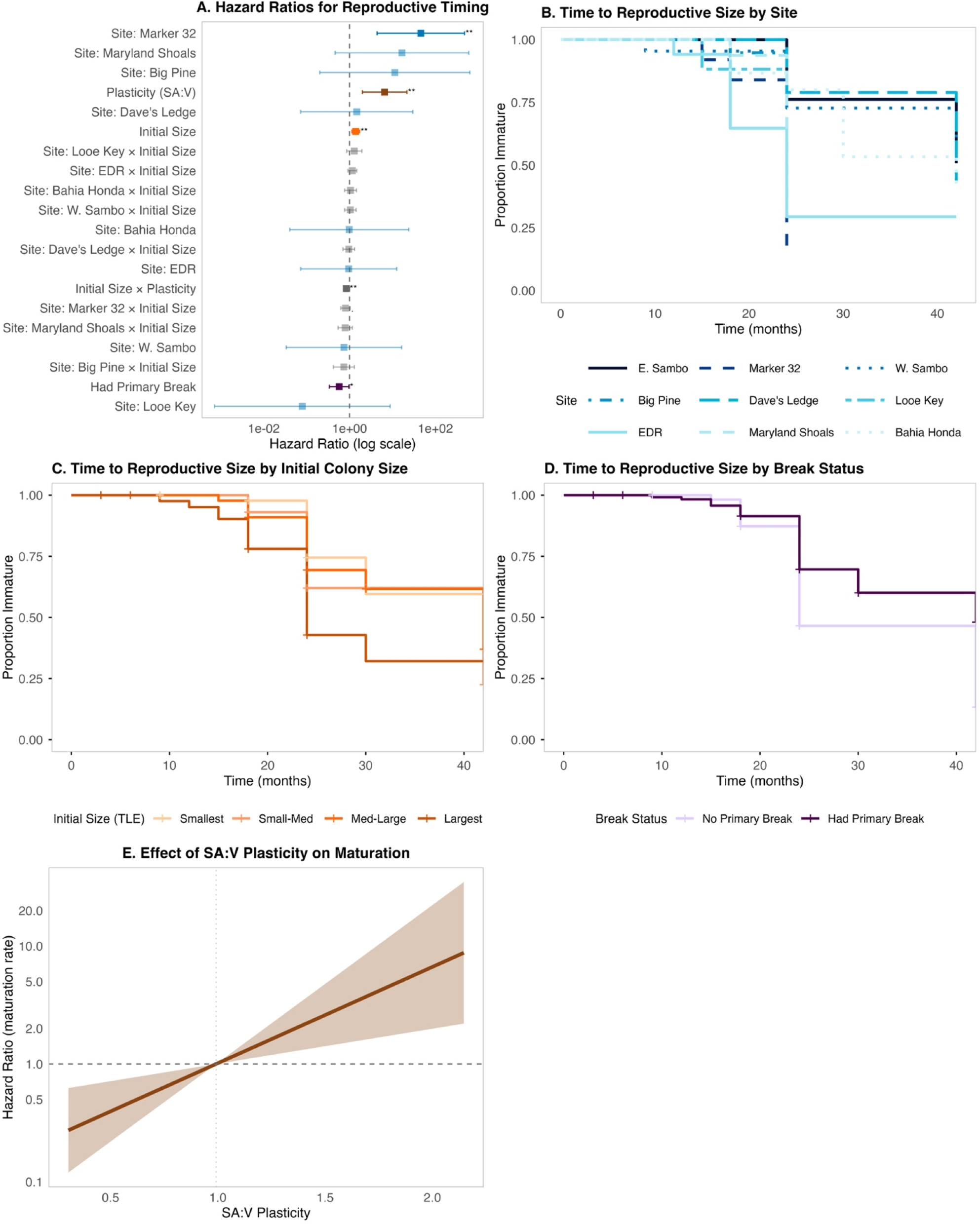
Multi-panel analysis of factors influencing time to sexual maturity in corals. **Panel A** displays hazard ratios and 95% confidence intervals from a Cox proportional hazards model examining effects of site (with Eastern Sambo as reference), initial colony size at transplantation, presence of primary branch breakage, and morphological plasticity (measured as surface area to volume ratio response). Points are color-coded by variable type: blue for sites, orange for initial size, dark purple for primary break status, brown for plasticity, and grey for interaction terms. Non-significant effects (p≥0.05) are shown with transparency. Points to the right of the vertical dashed line (HR>1) indicate factors associated with accelerated maturation, while those to the left (HR<1) indicate delayed maturation. Statistical significance is denoted by asterisks (* p<0.05, ** p<0.01, *** p<0.001). **Panel B** shows Kaplan-Meier curves for maturation by site (blue gradient), revealing substantial variation in maturation timing across reef locations. **Panel C** illustrates the effect of initial colony size (orange gradient), with larger corals (darkest orange) reaching maturity more quickly than smaller ones (lightest orange). **Panel D** demonstrates that primary branch breakage (dark purple) significantly delays sexual maturity compared to corals without primary breaks (light purple). **Panel E** shows the continuous effect of morphological plasticity (SA:V ratio) on sexual maturation hazard, revealing a strong positive relationship (HR = 6.56, p = 0.002) where corals with higher surface area to volume ratios mature significantly faster. The brown line represents the hazard ratio across the observed plasticity range (0.30-2.15), with shaded 95% confidence intervals.

Across all predictors, hazard ratios greater than 1 were associated with accelerated maturation (e.g., larger initial size, higher morphological plasticity, Marker 32 site), while hazard ratios less than 1 indicated delayed maturation (e.g., primary branch breakage). Physical damage, morphological traits, and environmental variation thus jointly influenced the timing of reaching reproductive size in *A. cervicornis*.

To determine whether plasticity aids recovery after damage rather than preventing damage, we compared breakage rates and post-breakage maturation between plasticity groups. High- and low-plasticity corals experienced similar rates of primary breaks (75.2% vs. 75.8%, χ^2^ test p = 0.734), indicating that plasticity does not confer resistance to mechanical damage. However, among the 119 corals that experienced primary breaks, high-plasticity individuals were significantly more likely to reach sexual maturity (44.0%, 22/50) than low-plasticity individuals (27.5%, 19/69; HR = 0.515 for low vs. high plasticity, p = 0.040). This demonstrates that plasticity specifically enhances post-breakage recovery and reproductive development.

#### Expected Reproductive Output

Von Bertalanffy growth curves fit to individual-level TLE data yielded asymptotic sizes of *L∞* =200 *cm* (*k* == 0.016, L_0_ = 6.0) for colonies without primary breaks and *L∞* = 150 *cm* (*k* == 0.010, *L*_*0*_ = 6.4) for colonies with primary breaks. By month 36, projected TLE was 89 cm (no primary break) versus 48 cm (primary break), corresponding to 56% and 30% of the Boisvert-converted reproductive threshold (∼161 cm TLE), respectively.

The demographic model estimated that colonies experiencing primary breaks produced 58% fewer expected oocytes over three spawning seasons compared with unfragmented colonies (Fig. 5). This reduction was driven by the combined effects of smaller colony volume (and therefore less gametogenic tissue) and lower survival probability at each spawn event. The result was robust to variation in all model parameters: across the full range of published fecundity values, von Bertalanffy asymptotes *L∞* = 100 - 300 *cm*, and logistic threshold assumptions, the estimated reduction remained between 56% and 76%. Notably, polyp density and oocytes per polyp did not affect the relative reduction, as these parameters scale in both groups equally.

**Figure 5.**
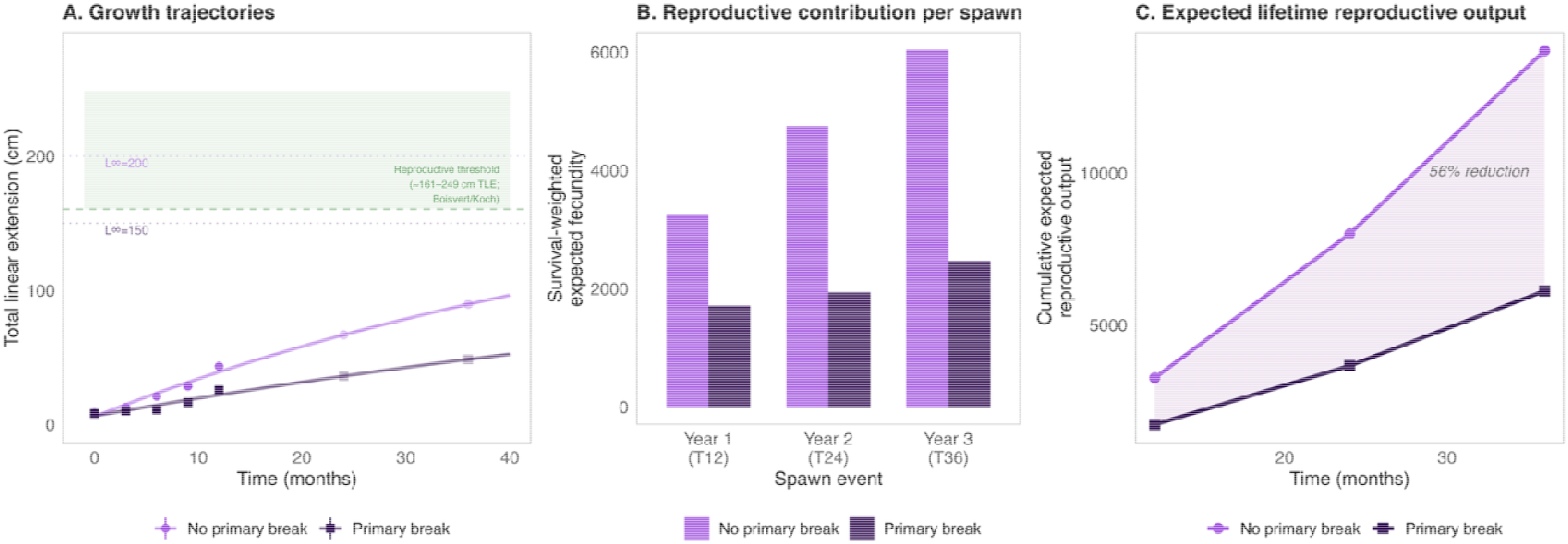
Expected reproductive output model comparing fragmented and unfragmented colonies over three spawning seasons. (A) Von Bertalanffy growth trajectories fit to individual-level TLE data from T0–T12 for colonies without primary breaks (medium purple; L∞ = 200 cm) and colonies with primary breaks (dark purple; L∞ = 150 cm). Solid points with standard error bars show observed mean TLE at each monitoring timepoint; open points show projected TLE at subsequent annual spawn events (T24, T36). The green shaded band indicates the reproductive size threshold (∼161–249 cm TLE) derived by converting Koch et al.’s (2022) ellipsoid volume threshold (∼5,000–10,000 cm^3^) to TLE using the regression from Kiel et al. (2012; Equation 2 in Boisvert et al. 2024). By month 36, unfragmented colonies reached 56% of the lower reproductive threshold compared to 30% for fragmented colonies. (B) Survival-weighted expected fecundity at each of three annual spawn events (T12, T24, T36), calculated as the product of Kaplan-Meier survival probability, colony volume estimated from the TLE-to-volume power-law fit to photogrammetry data, polyp density (16 polyps/cm^2^), and oocytes per polyp (7 oocytes/polyp), with fecundity parameters from Koch et al. (2022). (C) Cumulative expected reproductive output summed across all three spawn events. Colonies experiencing primary breaks produced an estimated 58% fewer expected oocytes than unfragmented colonies, a reduction driven by both smaller colony volume and lower survival probability. This estimate was robust across the full range of published fecundity parameter values, von Bertalanffy asymptotes (L∞ = 100–300 cm), and threshold assumptions (sensitivity range: 56– 76% reduction).

## Discussion

Primary-branch fragmentation delivers a double blow to staghorn corals: it more than doubles mortality risk and slows the pace of reaching a sexually mature size class by roughly 40%, resulting in an estimated 58% reduction in expected reproductive output (Fig. 5). Yet that delay is not inevitable. Colonies with high SA:V plasticity reached the reproductive size threshold significantly faster. This advantage likely emerges from damage-regrowth interactions that enable architectural reorganization in plastic genotypes. Breakage and plasticity vary sharply among genotypes (e.g. 41 and 13 break most often) and across sites (Bahia Honda, Looe Key, EDR show the highest risks), so the payoff of potential asexual propagation via primary-branch fragmentation depends on both heredity in the broad sense and habitat. In short, how a coral breaks—and how it regrows—jointly determine whether fragmentation becomes a demographic dead-end or a viable life-history tactic in the Anthropocene.

### The Reproductive Trade-off and Double Negative Effect of Breakage

Our results provide empirical evidence for a reproductive trade-off in *A. cervicornis*, where asexual reproduction through fragmentation undermines sexual reproductive potential by reducing colony size below critical thresholds required for maturation. Our survival analysis revealed that primary branch breaks significantly increased mortality risk of the original outplant. Importantly, the same breakage type also delayed sexual maturity in these damaged colonies which survived. This dual effect of primary breakage on the original colonies—impairing both survival and reproductive capacity—represents a “double negative” on local population dynamics that builds on prior observations in other *Acropora* species showing that gamete production may cease in fragmented colonies and their derivatives several years after major disturbance events such as hurricanes (Lirman, 1998). This pattern parallels size-dependent mortality-reproduction trade-offs documented across species, where damage-induced size reductions cascade through multiple fitness components (Stearns 1992; Madin et al. 2020).

We advance understanding of this phenomenon in several ways. First, we quantify reproductive delay magnitude using hazard ratios in a time-to-event framework rather than binary gamete presence. Second, we identify that primary branch breaks, not all fragmentation types, drive this reproductive impact. Third, we demonstrate this relationship persists across multiple sites and environmental conditions, suggesting a fundamental biological trade-off rather than context-dependent response. Fourth, our demographic model shows that even without invoking any maturity threshold, fragmentation reduces expected cumulative reproductive output by ∼58% purely through smaller colony volume and lower survival — an estimate that is conservative because it assumes no further breakage events during the growth projection. Using the Boisvert et al. (2024) conversion to express Koch et al.’s (2022) reproductive threshold in TLE, unfragmented colonies grew to 56% of the reproductive threshold TLE by month 36 (89 of ∼161 cm), while fragmented colonies grew to only 30% (48 of ∼161 cm), indicating that fragmentation substantially delays the approach to reproductive competence regardless of the threshold framework employed. However, the ecological dynamics of fragmentation are complex as the broken fragments themselves represent asexual propagules that may establish in new locations. While our analysis focused on the fate of the original outplants, the broader population consequences of fragmentation likely involve a balance between the documented negative effects on source colonies and the potential for successful fragment recruitment and establishment elsewhere (Highsmith, 1982; Garrison & Ward, 2012).

Previous analyses of survival and growth patterns over the initial twelve months did not find that fragmentation impacted survival (Million et al. 2022). Our contrasting result may be due to a longer observation period in the current analysis, or the decision to parse breakage by type. Lirman (2000) showed that fragmentation can severely impact colony fitness in Caribbean staghorn corals by reducing reproductive potential for up to 4 years post-fragmentation, with both the parent colonies and fragments showing complete absence of gametes during this recovery period. Darling et al. (2012) also highlighted the elevated mortality risks associated with size reductions from breakage in reef-building corals such as *Acropora cervicornis*. This aligns with broader patterns in reef corals where partial mortality acts disproportionately on smaller colonies, and size-dependent mortality creates fundamental constraints on colony fitness (Madin et al. 2020). Our findings extend these results by quantitatively linking breakage not only to increased mortality but also to delays in sexual maturity, underscoring its potential to disrupt long-term population recovery. Importantly, we found that morphological plasticity in surface area to volume ratio can mitigate these negative effects, with high-plasticity colonies reaching reproductive size ∼6× faster than low-plasticity counterparts (detailed analysis in Compensatory Role of Morphological Plasticity below).

Our findings align with the evolutionary history of Caribbean corals described by Roff (2021), who conceptualized how species like *A. cervicornis* may have evolved to rely primarily on asexual fragmentation rather than sexual reproduction. Tunnicliffe (1981) provided the first empirical evidence for this adaptation, documenting that over 80% of corals in Jamaican populations were broken from their bases yet had successfully re-anchored to grow rapidly, with little evidence of sexual recruitment. Historically, these corals maintained population sizes through rapid growth rates (10-15 cm/year) documented by Gladfelter et al. (1978) and Lewis (1974), enabling them to thrive despite high rates of fragmentation. The evolutionary significance of this strategy was further established by Highsmith (1982), who postulated that corals had evolved morphological and mechanical properties specifically facilitating fragmentation. However, the ‘double negative’ impact we observed suggests that modern environmental stressors have disrupted this reproductive mechanism. While fragmentation was once an adaptive strategy for Caribbean acroporids, our results indicate that under current conditions, it may instead contribute to population decline by simultaneously increasing mortality and delaying sexual maturity.

While Rapuano et al. (2023) demonstrated that post-pubescent corals may retain per-polyp reproductive capacity even when fragmented below species-specific size thresholds; whole-colony reproductive output still scales with colony size because larger colonies have proportionally more gametogenic tissue (Hall & Hughes 1996; Álvarez-Noriega et al. 2016; Koch et al. 2022). Moreover, the most phylogenetically relevant fragmentation experiments on branching *Acropora* (Okubo et al. 2007) found that small fragments of *A. formosa* resorbed their oocytes entirely, suggesting that retention of reproductive capacity in post-pubescent fragments (Rapuano et al. 2023) may not be generalizable across acroporids and warrants further study.

This interplay between breakage susceptibility, morphological plasticity, and reproductive timing reveals a complex adaptive landscape in which selection may favor genotypes that not only resist catastrophic breakage but also leverage plasticity to optimize fitness under variable environmental conditions. The pronounced genotypic differences we observed in breakage susceptibility—with some genotypes experiencing catastrophic breakage in nearly all colonies (e.g., Genotype 13), while others showed markedly lower rates—suggest that natural selection could act on this trait. This introduces an evolutionary dimension to fragmentation that has not been previously explored in coral demographic studies.

### Genotypic and Environmental Influences on Breakage

Genotype had a profound effect on breakage susceptibility. Notably, Genotype 13 exhibited 100% breakage rates with a significantly elevated hazard ratio for recurrent breaks, while Genotype 41 showed similarly high vulnerability. In contrast, Genotype 36 demonstrated greater resistance to breakage, serving as the reference genotype in our analyses. These patterns underscore that fragmentation potential varies among genets of *A. cervicornis* (Million et al. 2022) and may have a heritable component, suggesting that natural selection could act on breakage susceptibility in response to environmental pressures. This genotypic variation in breakage susceptibility likely reflects differences in colony architecture, as mechanical vulnerability to hydrodynamic dislodgement is strongly determined by growth form in corals (Madin et al. 2014). Such architecture-environment interactions influencing damage susceptibility may represent a general phenomenon in sessile organisms with complex morphologies. Given that fragments used in this experiment were clonal replicates of each genotype, the observed variation in breakage susceptibility among genotypes indicates a significant genetic component to this trait. The coral literature has evolved significantly in this understanding—early foundational studies documented species-level patterns (Tunnicliffe 1981; Gilmore & Hall 1976; Shinn 1976), with Highsmith (1982) first proposing that fragmentation might represent an adaptive life-history strategy. Connell (1973) established size-based mortality patterns in corals, which later supported the hypothesis that fragmentation would be adaptive primarily in species with low mortality rates above certain size thresholds. Our finding challenges previous assertions that fragmentation results primarily from within-colony variation or environmental factors with no genetic basis (Highsmith 1982) and provides empirical evidence that fragmentation susceptibility itself may be heritable and thus subject to evolutionary processes.

Site also dramatically affected breakage patterns, with locations such as Bahia Honda showing nearly universal breakage (96.6% of corals) and significantly higher hazard for recurrent breaks. Similarly, Looe Key and EDR demonstrated elevated break risk. Notably, both Looe Key and EDR are major tourist destinations, suggesting that human activity may be a significant contributor to the elevated breakage rates observed at these sites. This highlights how anthropogenic factors, particularly tourism pressure, may interact with natural environmental conditions to increase mechanical stress on corals. The pronounced site effects suggest that local environmental factors, including wave exposure, current patterns, and human activities, significantly influence coral mechanical vulnerability.

Finally, our temporal analysis revealed a pattern in breakage susceptibility over time, with rates initially increasing after outplanting, then decreasing, before increasing again as colonies matured. This non-linear relationship suggests that breakage risk may be tied to developmental stages. As colonies first establish, they may be particularly vulnerable. The subsequent decrease in breakage likely reflects successful establishment, while the later increase in breakage rates for mature colonies indicates that as corals grow larger, they may reach a structural threshold where their branching architecture again increases vulnerability to mechanical stress.

### Compensatory Role of Morphological Plasticity

Our time-to-event analysis for sexual maturity revealed that morphological plasticity in SA:V ratio serves as a critical compensatory mechanism following fragmentation damage. Higher SA:V plasticity significantly accelerated time to achieving a sexually mature size class. This acceleration suggests that corals with greater morphological flexibility may better acclimatize to local conditions and reach reproductive maturity sooner despite the costs of fragmentation.

This moderating effect identifies a mechanism through which corals balance competing demands of regrowth and reproductive output. High SA:V plasticity could enable colonies to maximize light capture and nutrient exchange while minimizing structural vulnerability (Enríquez et al. 2005; Madin 2005; Madin 2020). This morphological flexibility could allow colonies to allocate resources more efficiently between tissue repair and growth, reaching the size threshold for reproduction despite the energetic costs of fragmentation damage. Previous work has proposed that fragmentation disrupts reproduction by shifting energy allocation toward stabilization, lesion repair, and regrowth (Lirman 2000). Our findings support this resource allocation hypothesis and extend it by demonstrating that morphological plasticity may optimize how resources are distributed, potentially enabling colonies to invest simultaneously in both recovery and reproductive development. This represents a novel mechanism for plasticity-mediated mitigation of reproductive trade-offs. Our findings provide empirical support for theoretical predictions that plasticity can buffer demographic costs of environmental variability (Chevin et al. 2013), extending this framework to show that plasticity specifically compensates for mechanical damage rather than preventing it.

Importantly, because this plasticity is genotype-specific, this mechanism may be subject to selection, adding an evolutionary dimension to its role in shaping reproductive trajectories. The significant interaction between initial size and plasticity further refines our understanding of how these factors work together to influence reproductive timing. Our stratified analysis reveals that morphological plasticity enables exploitation of certain breakage events, possibly for beneficial architectural reorganization. Remarkably, corals with secondary breaks showed lower mortality than unbroken colonies regardless of plasticity level (0% vs. 10.5% in high-plasticity; 7.7% vs. 20.8% in low-plasticity), though the effect was strongest in high-plasticity individuals. Moreover, high-plasticity corals’ superior maturation success following primary breaks (44% vs. 27.5%) reveals that plasticity may also enhance regenerative capacity, likely through optimized regrowth patterns and resource reallocation during recovery. This suggests the compensatory mechanism yields greater fitness benefit to high-plasticity individuals. These results help reconcile the apparent paradox of fragmentation as both an asexual reproductive strategy and a mortality risk. For high-plasticity genotypes experiencing moderate damage, fragmentation may function as an adaptive response through enhanced regrowth patterns that optimize colony architecture. However, severe damage (primary breaks) overwhelms any compensatory capacity regardless of plasticity level, indicating that the location and severity of damage critically determine whether fragmentation becomes detrimental or advantageous. This compensatory role may explain the evolution of fragmentation as a life-history strategy in disturbed environments where mechanical damage is unavoidable (Roff 2021). Future work should aim to investigate whether this mechanism operates in other modular organisms where fragmentation interacts with size-dependent reproduction, including bryozoans, colonial ascidians, and clonal plants (Hughes 1989; Powell & Burgess 2025).

### Broader Context and Implications for Restoration

Our findings underscore the importance of quantifying individual colony fate. The genotypic basis of breakage susceptibility adds an important dimension to coral life history, suggesting the balance between sexual and asexual reproduction is subject to selection. Morphological plasticity counteracting fragmentation-induced reproductive delays identifies a mechanism through which plasticity enhances fitness beyond immediate survival, linking genotype-specific morphological responses to their demographic consequences (Smallegange et al. 2025).

These insights directly apply to coral reef restoration, which routinely propagates and outplants fragments across diverse reef sites (Schopmeyer et al. 2017). Identifying genotypes with lower breakage risk and higher plasticity could enable practitioners to select coral stocks better equipped to maintain balanced reproductive strategies, enhancing long-term recovery. Our results suggest genotypes like 36 (showing greater breakage resistance) combined with sites promoting faster maturation (like Marker 32) offer optimal restoration conditions. Accounting for genotypic breakage susceptibility and its reproductive implications could potentially double reproductive output in restored populations, dramatically improving recovery timelines. As warming oceans intensify bleaching frequency and severity(Hughes et al. 2017; Manzello et al. 2025), our findings provide crucial guidance for propagating genotypes maintaining balanced reproductive strategies under increasing stress.

## Acknowledgements

We are grateful to Yingqi Zhang, Hunter Ramo, Cory Walter, and Joseph Kuehl for their help in photographing coral in situ. ACV was funded by a University of Southern California Presidential Sustainability Fellowship, which also funded CH and DZ. All fieldwork was conducted under permits FKNMS-2015-163-A1 and FKNMS-2018-035. This research was supported by the National Oceanic and Atmospheric Administration Coral Reef Conservation Program grant NA17NOS4820084, the National Science Foundation IOS-2222272, and private funding from the Alfred P. Sloan and Rose Hills Foundations to CDK.

