## Supplement for "Reproductive Trade-offs in Caribbean Corals: Plasticity Accelerates While Fragmentation Delays Reproductive Capacity"

### 1. EXPERIMENTAL DESIGN AND METHODS

This study extends the 12-month dataset originally described in Million et al. (2022) by reanalyzing their data and extending the monitoring period to a maximum of 42 months to capture additional growth and reproductive milestones. The experiment involved ten distinct coral genotypes of *Acropora cervicornis* sourced from the lower Florida Keys and maintained at Mote Marine Laboratory's in situ coral nursery prior to outplanting. In April 2018, 270 coral ramets (27 per genotype, with a mean Total Linear Extension (TLE) of 8.4 cm) were outplanted to nine active restoration sites across the Florida Keys reef tract at depths ranging from 5.6 to 9.1 m. At each site, three ramets per genotype were randomly allocated to each of three ten-coral arrays. Ramets were photographed immediately before transplantation and measured for TLE, then re-surveyed quarterly during the first year and biannually for an additional 2.5 years.

### 2. BREAKAGE PATTERNS

#### 2.1 Overall Breakage Distribution

Detailed breakdown of coral fragmentation events by severity and location.

#### Table S1: Break Pattern Distribution

| **Break Type** | **Count** | **Percentage** |
| --- | --- | --- |
| Primary, Catastrophic | 177 | 67.3% |
| Primary, Partial | 18 | 6.8% |
| Secondary, Catastrophic | 13 | 4.9% |
| Secondary, Partial | 3 | 1.1% |
| Tertiary, Catastrophic | 5 | 1.9% |
| None (no breaks) | 42 | 16.0% |
| NA (break but unknown type) | 5 | 1.9% |

Analysis of 263 unique corals revealed that 83.7% (220 corals) experienced at least one breakage event during the study period. The most common fragmentation type was Primary Catastrophic (1C), accounting for 67.3% of all breaks, while 16.0% of corals remained intact throughout the observation period.

#### 2.2 Genotypic Variation in Breakage

#### Table S2: Genotype-Specific Break Patterns

Details of genotypic variation in susceptibility to breakage.

**Genotype Break Analysis**

| **Genotype** | **Total Corals** | **Corals With Breaks** | **Percent With Breaks** | **Avg Breaks Per Colony** | **Corals With Classified Break Type** | **Percent Corals With Classified Break Type** |
| --- | --- | --- | --- | --- | --- | --- |
| 36 | 27 | 20 | 74.1% | 1.04 | 20 | 100% |
| 1 | 26 | 20 | 76.9% | 1.31 | 20 | 100% |
| 50 | 26 | 19 | 73.1% | 1.42 | 19 | 100% |
| 3 | 27 | 24 | 88.9% | 1.37 | 24 | 100% |
| 44 | 27 | 22 | 81.5% | 1.41 | 22 | 100% |
| 7 | 26 | 23 | 88.5% | 1.31 | 23 | 100% |
| 31 | 26 | 22 | 84.6% | 1.35 | 22 | 100% |
| 13 | 27 | 27 | 100% | 1.37 | 25 | 92.6% |
| 62 | 25 | 19 | 76% | 1.00 | 17 | 89.5% |
| 41 | 26 | 24 | 92.3% | 1.31 | 23 | 95.8% |

Breakage patterns varied substantially among genotypes, ranging from 73.1% to 100% of colonies experiencing fragmentation. Genotype 13 had the highest break rate with all 27 colonies (100%) experiencing breaks, followed by Genotype 41 (92.3%) and Genotypes 3 and 7 (both approximately 89%). Genotypes 50 and 36 showed the lowest breakage rates (73.1% and 74.1%, respectively). These pronounced differences in breakage susceptibility across genotypes, despite each genotype being replicated across all sites, suggest that fragmentation susceptibility has a heritable component.

#### 2.3 Site Effects on Breakage

#### Table S3: Site-Specific Break Patterns

Break occurrence across different study sites.

| **Site** | **Total Corals** | **Corals With Breaks** | **Corals With Break Type Info** | **Percent With Break Type Info** |
| --- | --- | --- | --- | --- |
| E. Sambo | 30 | 20 | 20 | 100% |
| Marker 32 | 30 | 20 | 20 | 100% |
| W. Sambo | 30 | 22 | 22 | 100% |
| Big Pine | 28 | 25 | 25 | 100% |
| Dave's Ledge | 28 | 27 | 26 | 96.3% |
| Looe Key | 29 | 27 | 27 | 100% |
| EDR | 29 | 25 | 23 | 92% |
| Maryland Shoals | 30 | 26 | 26 | 100% |
| Bahia Honda | 29 | 28 | 26 | 92.9% |

Break rates varied substantially across sites, ranging from 66.7% to 96.6%. Bahia Honda and Dave's Ledge showed the highest fragmentation rates (96.6% and 96.4%, respectively), while E. Sambo and Marker 32 had the lowest (both 66.7%). Notable intermediate-to-high break rates occurred at tourist-frequented sites including Looe Key (93.1%) and EDR (86.2%), suggesting that human activities may contribute to mechanical stress on corals alongside natural environmental factors such as wave exposure and current patterns.

#### 2.4 Temporal Patterns of Breakage

#### Figure S1: Percentage of Corals with Breaks by Genotype

Bar chart showing the proportion of coral colonies experiencing at least one breakage event across ten genotypes of *A. cervicornis*. Genotypes ordered by break susceptibility (range: 73.1-100%). Error bars represent ±1 SE.

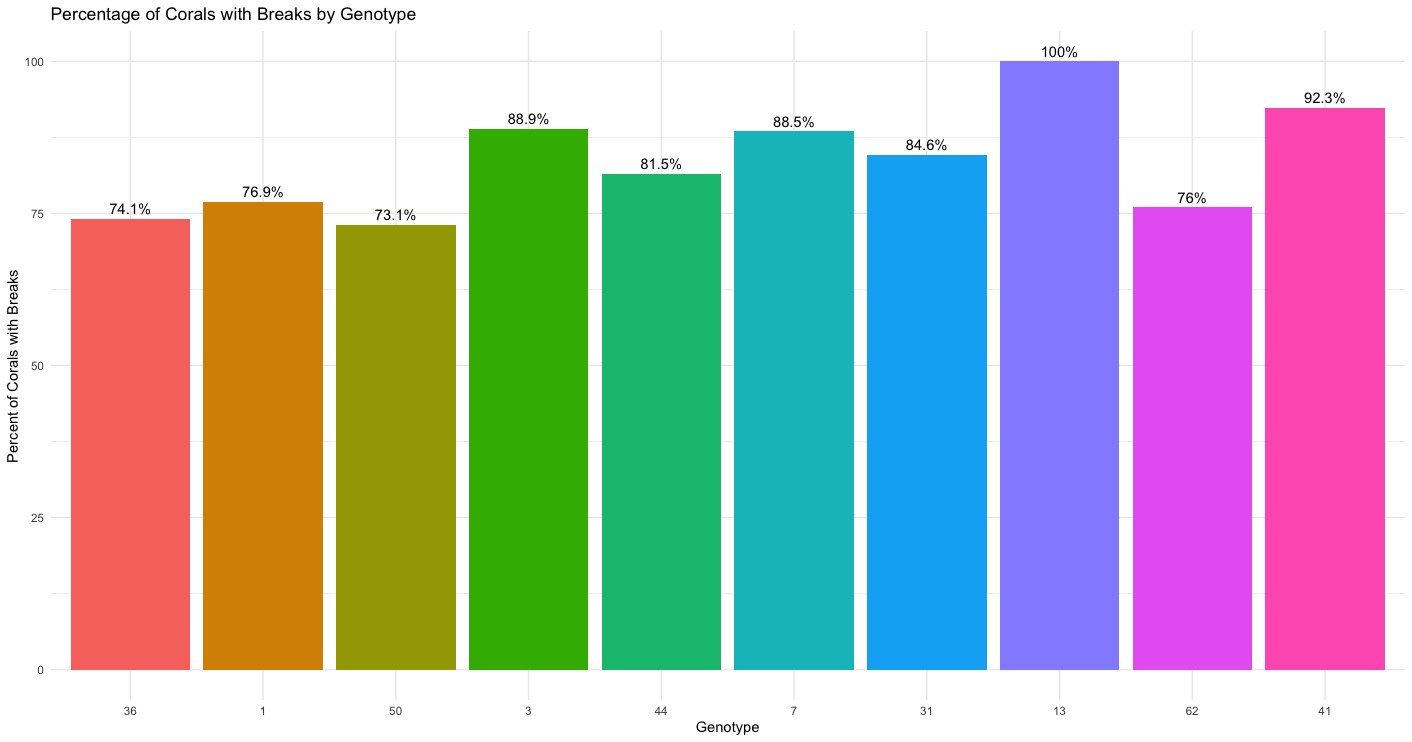

#### Figure S2: Distribution of Most Severe Break Type by Genotype.

Stacked bar plot showing the proportional distribution of break types by genotype. Break types categorized by severity: Primary Catastrophic (1C), Primary Partial (1P), Secondary Catastrophic (2C), Secondary Partial (2P), Tertiary Catastrophic (3C), and No Break. Genotypes ordered by increasing break susceptibility.

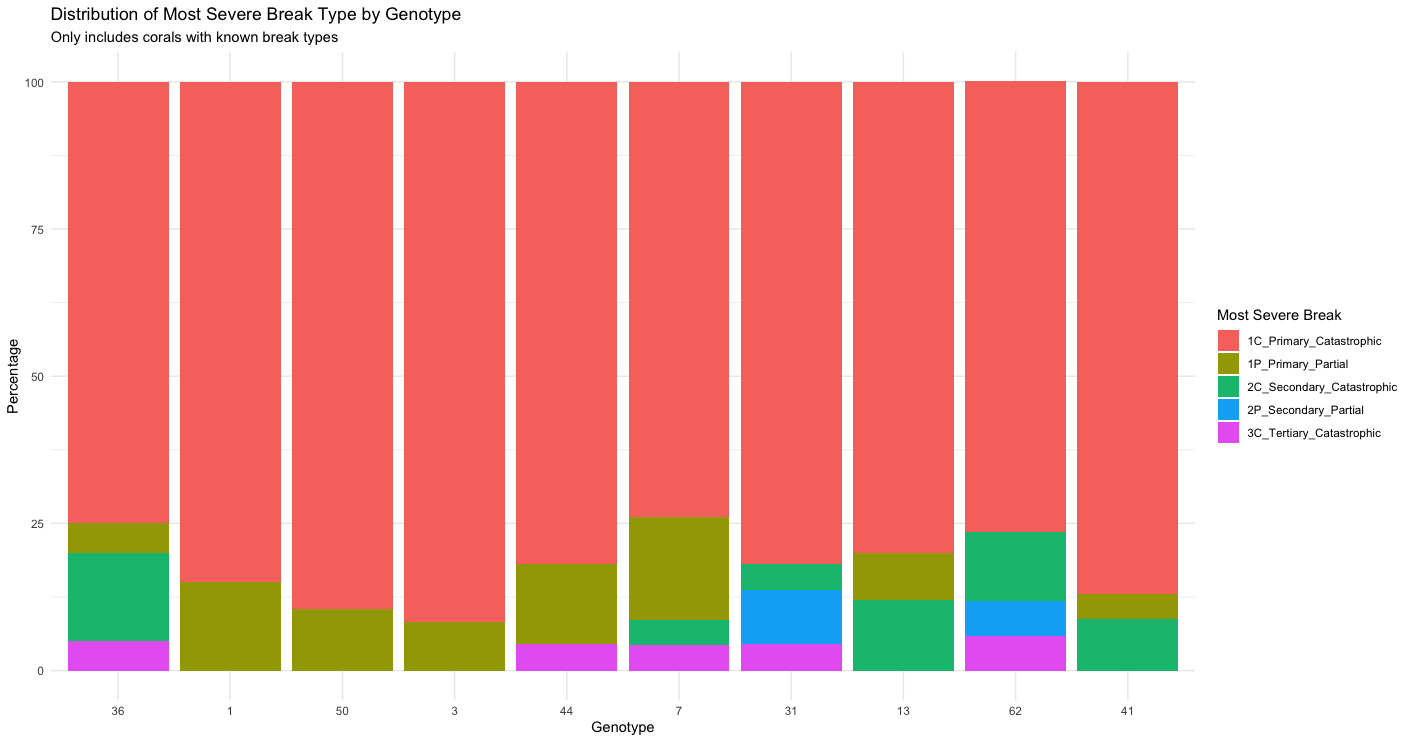

### 3. SURVIVAL ANALYSES

#### 3.1 Cox Proportional Hazards Models

#### Table S4: Cox Proportional Hazards Models for Coral Survival.

This table presents the results of Cox proportional hazards analyses examining factors affecting coral mortality, showing hazard ratios with 95% confidence intervals and p-values for site, genotype, and break type effects. The model can be expressed as h(t) = h₀(t) × exp(β₁Site + β₂Genotype), where h(t) is the hazard function representing the risk of mortality at time t, h₀(t) is the baseline hazard function, and β₁ and β₂ are the estimated coefficients for site and genotype effects, respectively.

##### Site and Genotype Effects (compared to reference site/genotype)

| **Site/Genotype** | **Hazard Ratio** | **95% CI Lower** | **95% CI Upper** | **p-value** |
| --- | --- | --- | --- | --- |
| E. Sambo | 1.61 | 0.53 | 4.92 | 0.405 |
| W. Sambo | 1.69 | 0.55 | 5.19 | 0.355 |
| Big Pine | 2.82 | 0.99 | 8.03 | 0.051 |
| Dave's Ledge | 1.87 | 0.61 | 5.73 | 0.272 |
| Looe Key | 4.25 | 1.54 | 11.74 | **0.005** |
| EDR | 3.81 | 1.37 | 10.60 | **0.010** |
| Maryland Shoals | 3.19 | 1.15 | 8.86 | **0.026** |
| Bahia Honda | 5.00 | 1.82 | 13.71 | **0.002** |
| 36 | 1.71 | 0.57 | 5.10 | 0.337 |
| 1 | 2.93 | 1.04 | 8.24 | **0.041** |
| 3 | 1.83 | 0.61 | 5.46 | 0.279 |
| 44 | 1.85 | 0.62 | 5.52 | 0.270 |
| 7 | 2.11 | 0.72 | 6.19 | 0.174 |
| 31 | 2.22 | 0.76 | 6.51 | 0.146 |
| 13 | 2.99 | 1.05 | 8.51 | **0.040** |
| 62 | 3.44 | 1.19 | 9.96 | **0.022** |
| 41 | 3.36 | 1.18 | 9.56 | **0.023** |

#### 3.2 Model Validation and Diagnostics

#### **Table S5: Proportional Hazards Assumption Tests for Cox Model with Break Type.**

Tests confirm the proportional hazards assumption is satisfied for the Cox model including break type (all p > 0.05). Simplified Break Type categorizes colonies as: No Break (reference), Primary_Branch (1C+1P), or Secondary_or_Other (2C+2P+3C+4C)

#### **Proportional Hazards Test for Model Including Break Type**

| **Variable** | **Chi-square** | **df** | **p-value** |
| --- | --- | --- | --- |
| Site | 5.05 | 8 | 0.752 |
| Genotype | 15.95 | 9 | 0.068 |
| Simplified Break Type | 0.60 | 2 | 0.741 |
| GLOBAL | 27.05 | 19 | 0.104 |

In this model, all variables satisfy the proportional hazards assumption (p > 0.05), suggesting that adding break type improves the model's validity.

#### 3.3 Effects of Breakage on Mortality Risk

#### Table S6: Cox Proportional Hazards Model with Break Type.

##### This table presents the results of an enhanced Cox proportional hazards model that incorporates break type as an additional predictor variable. The model can be expressed as h(t) = h₀(t) × exp(β₁Site + β₂Genotype + β₃BreakType), where h(t) is the hazard function representing the risk of mortality at time t, h₀(t) is the baseline hazard function, and β₁, β₂, and β₃ are the estimated coefficients for site, genotype, and break type effects, respectively. The inclusion of break type improves model validity as confirmed by the proportional hazards tests.

##### Site, Genotype, and Break Type Effects (compared to reference site)

| **Site/Genotype/Break** | **Hazard Ratio** | **95% CI Lower** | **95% CI Upper** | **p-value** |
| --- | --- | --- | --- | --- |
| E. Sambo | 1.54 | 0.50 | 4.72 | 0.450 |
| W. Sambo | 1.84 | 0.60 | 5.67 | 0.285 |
| Big Pine | 2.13 | 0.74 | 6.14 | 0.162 |
| Dave's Ledge | 1.64 | 0.53 | 5.10 | 0.394 |
| Looe Key | 3.02 | 1.08 | 8.45 | **0.035** |
| EDR | 2.98 | 1.03 | 8.60 | **0.043** |
| Maryland Shoals | 2.81 | 1.00 | 7.88 | **0.050** |
| Bahia Honda | 3.20 | 1.13 | 9.07 | **0.029** |
| 36 | 1.90 | 0.63 | 5.72 | 0.251 |
| 1 | 2.89 | 1.03 | 8.13 | **0.045** |
| 3 | 1.60 | 0.54 | 4.81 | 0.399 |
| 44 | 1.79 | 0.60 | 5.36 | 0.301 |
| 7 | 2.11 | 0.72 | 6.20 | 0.175 |
| 31 | 2.46 | 0.83 | 7.25 | 0.103 |
| 13 | 2.47 | 0.83 | 7.29 | 0.103 |
| 62 | 3.87 | 1.31 | 11.44 | **0.014** |
| 41 | 2.97 | 1.02 | 8.63 | **0.045** |
| Primary Branch | 2.58 | 1.14 | 5.82 | **0.022** |
| Secondary Branch or Other | 0.20 | 0.02 | 1.70 | 0.142 |

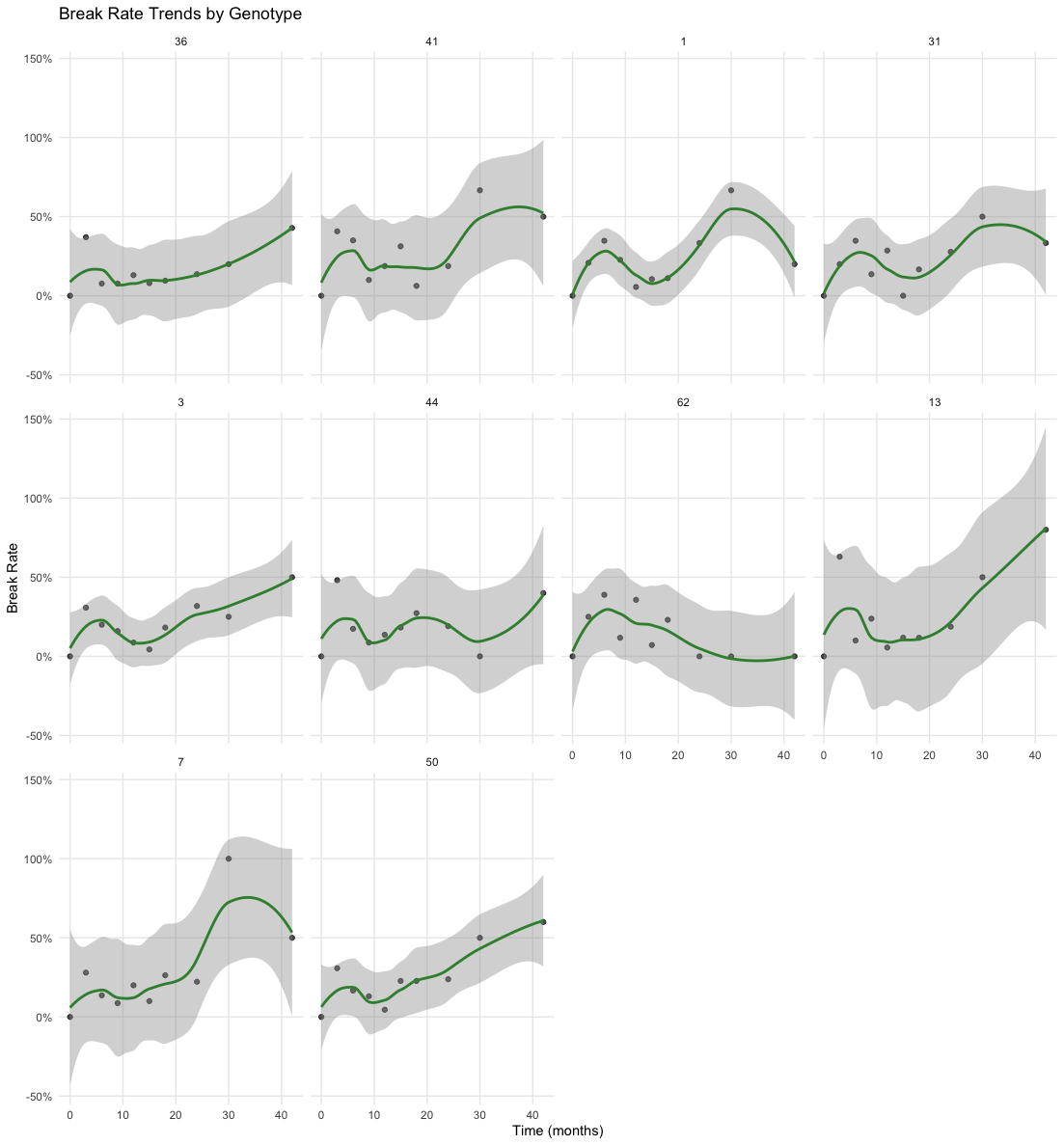

#### Figure S3: Break Rate Trends by Genotype Over Time.

This figure shows the temporal changes in coral breakage probability across different genotypes. Each colored line represents the smoothed trend line for that genet, calculated using LOESS regression. The shaded area surrounding each line represents the 95% confidence interval of the estimated trend, indicating the uncertainty in the predicted breakage probability. Data points show the observed break rates at each time point. Break rates are presented as percentages at each given time-point, with higher values indicating greater susceptibility to breakage.

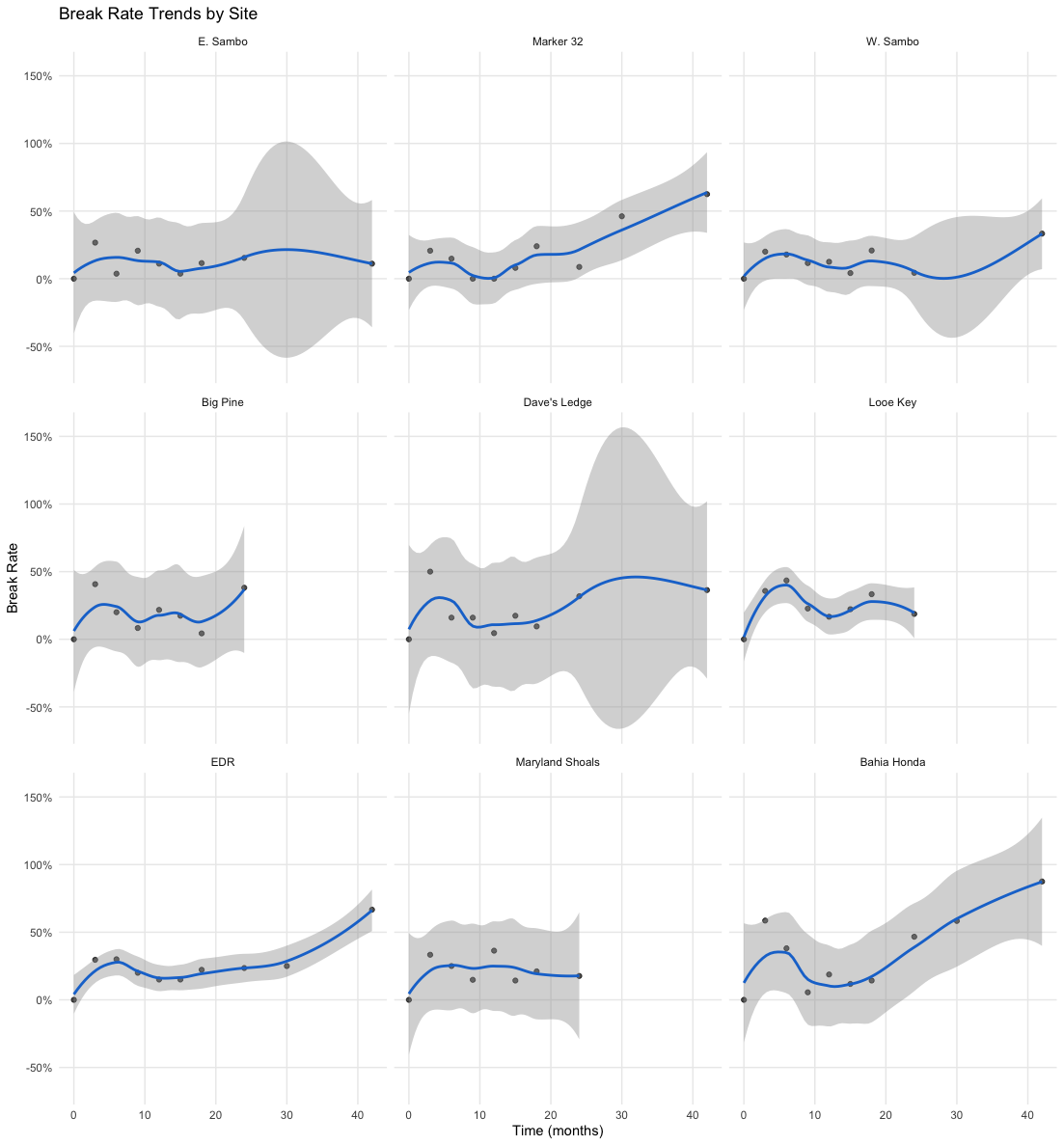

#### Figure S4: Break Rate Trends by Site Over Time.

This figure shows the temporal changes in coral breakage probability across different study sites. Each colored line represents the smoothed trend line for a different site, calculated using LOESS regression. The shaded area surrounding each line represents the 95% confidence interval of the estimated trend, indicating the uncertainty in the predicted breakage probability. Data points show the observed break rates at each time point. Break rates are presented as percentages at each given time-point, with higher values indicating greater susceptibility to breakage.

#### Table S7: Logistic Regression Analysis of Factors Affecting Breakage Probability

Binary logistic regression examining the effects of genotype, site, and initial size (T0_TLE) on breakage occurrence. The model can be expressed as:

logit(p) = log(p/(1-p)) = β₀ + β₁Genotype + β₂Site + β₃T0_TLE

where p is the probability of a coral experiencing a break, β₀ is the intercept, and β₁, β₂, and β₃ are the estimated coefficients for genotype, site, and initial coral size (T0_TLE) effects, respectively.

| **Statistic** | **Value** |
| --- | --- |
| Null deviance | 1718.1 on 1799 degrees of freedom |
| Residual deviance | 1667.9 on 1781 degrees of freedom |
| AIC | 1705.9 |
| Fisher Scoring iterations | 4 |

#### Genotype, Site, and Initial Size Effect on Break Probability

| **Genotype/Site/Size** | **Estimate** | **Std. Error** | **z value** | **p-value** | **Significance** |
| --- | --- | --- | --- | --- | --- |
| 41 | 0.759 | 0.288 | 2.639 | 0.008 | ** |
| 1 | 0.442 | 0.294 | 1.506 | 0.132 |  |
| 31 | 0.451 | 0.287 | 1.571 | 0.116 |  |
| 3 | 0.435 | 0.279 | 1.557 | 0.119 |  |
| 44 | 0.508 | 0.276 | 1.840 | 0.066 | . |
| 62 | 0.526 | 0.311 | 1.690 | 0.091 | . |
| 13 | 0.585 | 0.281 | 2.084 | 0.037 | * |
| 7 | 0.428 | 0.286 | 1.495 | 0.135 |  |
| 50 | 0.514 | 0.279 | 1.845 | 0.065 | . |
| Marker 32 | 0.154 | 0.286 | 0.540 | 0.589 |  |
| W. Sambo | 0.092 | 0.294 | 0.314 | 0.753 |  |
| Big Pine | 0.548 | 0.279 | 1.964 | 0.050 | * |
| Dave's Ledge | 0.623 | 0.276 | 2.255 | 0.024 | * |
| Looe Key | 0.880 | 0.278 | 3.171 | 0.002 | ** |
| EDR | 0.726 | 0.282 | 2.576 | 0.010 | ** |
| Maryland Shoals | 0.696 | 0.276 | 2.520 | 0.012 | * |
| Bahia Honda | 1.295 | 0.266 | 4.861 | <0.001 | *** |
| T0_TLE (Initial Size) | -0.050 | 0.024 | -2.040 | 0.041 | * |

#### Table S8. Mortality rates by break type and morphological plasticity level.

Corals were classified as high or low plasticity based on median surface area to volume ratio plasticity (median = 0.87). Sample sizes and mortality outcomes are shown for each combination of plasticity level and break category.

| **Plasticity Group** | **Break Category** | **N** | **Mortality (n/N)** | **Mortality %** |
| --- | --- | --- | --- | --- |
| High Plasticity | No Break | 19 | 2/19 | 10.5% |
| High Plasticity | Primary Break | 79 | 32/79 | 40.5% |
| High Plasticity | Secondary/Other Break | 7 | 0/7 | 0% |
| Low Plasticity | No Break | 24 | 5/24 | 20.8% |
| Low Plasticity | Primary Break | 116 | 56/116 | 48.3% |
| Low Plasticity | Secondary/Other Break | 13 | 1/13 | 7.7% |

### 4. RECURRENT BREAKAGE ANALYSES

#### 4.1 Andersen-Gill Cox Model Overview

#### Table S9: Andersen-Gill Cox Model for Recurrent Breakage Events.

We employed the Andersen-Gill extension of the Cox proportional hazards model to analyze recurrent breakage events, accounting for within-coral correlation when colonies experienced multiple breaks over time. The model showed good fit (concordance = 0.625, SE = 0.015) and satisfied the proportional hazards assumption (p = 0.42). Table S10: Andersen-Gill Cox Model for Recurrent Breakage Events Analysis of 1,576 colony-time intervals capturing 339 breakage events.

Model: h(t|coral i) = h₀(t) × exp(β₁Genotype + β₂Site + β₃T0_TLE),

where h_i(t) is the hazard function for the ith coral at time t; h₀(t) is the baseline hazard function; and β₁, β₂, and β₃ are coefficients for genotype, site, and initial size effects. The Andersen-Gill approach allows analysis of multiple breakage events per colony.

#### **Model Overview**

| **Statistic** | **Value** |
| --- | --- |
| Sample size | 1,576 colony-time intervals |
| Number of events | 339 break events |
| Concordance | 0.625 (se = 0.015) |
| Global Wald test | χ²(18) = 240, p < 2e-16 |
| Likelihood ratio test | χ²(18) = 51.65, p = 4e-5 |
| Proportional hazards test | χ²(18) = 18.47, p = 0.42 (GLOBAL) |

#### 4.2 Genotype, Site, and Initial Size Effects on Recurrent Breakage

#### **Genotype Hazard Ratios for Recurrent Breakage**

| **Genotype/Site/Size** | **Hazard Ratio** | **95% CI Lower** | **95% CI Upper** | **z value** | **p-value** | **Significance** |
| --- | --- | --- | --- | --- | --- | --- |
| 1 | 1.53 | 1.00 | 2.34 | 1.945 | 0.052 | . |
| 50 | 1.54 | 0.97 | 2.44 | 1.830 | 0.067 | . |
| 3 | 1.49 | 0.96 | 2.30 | 1.784 | 0.074 | . |
| 44 | 1.62 | 1.05 | 2.50 | 2.174 | 0.030 | * |
| 7 | 1.46 | 0.90 | 2.35 | 1.541 | 0.123 |  |
| 31 | 1.52 | 1.02 | 2.25 | 2.080 | 0.038 | * |
| 13 | 1.78 | 1.27 | 2.50 | 3.337 | <0.001 | *** |
| 62 | 1.63 | 0.94 | 2.85 | 1.724 | 0.085 | . |
| 41 | 1.94 | 1.27 | 2.95 | 3.085 | 0.002 | ** |
| Marker 32 | 1.07 | 0.67 | 1.72 | 0.294 | 0.769 |  |
| W. Sambo | 1.08 | 0.68 | 1.71 | 0.321 | 0.748 |  |
| Big Pine | 1.82 | 1.16 | 2.88 | 2.580 | 0.010 | ** |
| Dave's Ledge | 1.74 | 1.17 | 2.59 | 2.729 | 0.006 | ** |
| Looe Key | 2.40 | 1.56 | 3.68 | 4.003 | <0.001 | *** |
| EDR | 1.90 | 1.37 | 2.63 | 3.855 | <0.001 | *** |
| Maryland Shoals | 1.95 | 1.13 | 3.39 | 2.390 | 0.017 | * |
| Bahia Honda | 2.93 | 1.85 | 4.63 | 4.594 | <0.001 | *** |
| T0_TLE (Initial Size) | 0.96 | 0.93 | 1.00 | -2.074 | 0.038 | * |

### 5. TIME TO SEXUAL MATURITY ANALYSES

Detailed results from Cox proportional hazards models examining factors affecting time to sexual maturity. We begin with an additive model.

#### 5.1 Model Selection Approach

##### Statistical Analysis – Model Selection for Time to Sexual Maturity

Cox proportional hazards models were fitted using the survival package in R (version 4.2.0) to analyze factors affecting time to sexual maturity. The initial model included all four-way interactions between Site, T0_TLE (initial size), HadPrimaryBreak, and Avg_Plas_SAtoV (morphological plasticity).

Model simplification proceeded through stepwise removal of interaction terms with p > 0.1, evaluated using likelihood ratio tests between nested models and changes in AIC. Terms representing biologically meaningful effects were retained regardless of statistical significance. The final model included main effects for all four predictors plus two interaction terms: T0_TLE × Avg_Plas_SAtoV (p = 0.006) and Site:Marker32 × T0_TLE (p = 0.086). Higher-order interactions and weaker two-way interactions were excluded after confirming their removal did not significantly reduce model performance based on likelihood ratio tests and AIC comparisons.

The optimized model demonstrated excellent discriminative ability (concordance = 0.813, SE = 0.026) and strong statistical significance (likelihood ratio test: χ²(20) = 72.68, p = 7e-08). All variables satisfied the proportional hazards assumption (p > 0.05), confirming model validity.

#### 5.2 Simple Additive Model Results

Initial analysis using a model with main effects only (n = 177 corals, 76 events) showed good fit (concordance = 0.777, SE = 0.031) and high statistical significance (LR test: χ²(11) = 56.85, p < 0.001).

#### Table S10: Cox Proportional Hazards Model for Time to Sexual Maturity - Additive Effects.

This table presents results from the simple additive model examining factors affecting time to sexual maturity in *Acropora cervicornis*. The model can be expressed as:

h(t) = h₀(t) × exp(β₁Avg_Plas_SAtoV + β₂HadPrimaryBreak + β₃T0_TLE + β₄Site)

where h(t) represents the hazard (rate) of reaching sexual maturity. Hazard ratios greater than 1 indicate factors accelerating maturation, while values less than 1 indicate factors delaying maturation.

#### **Hazard Ratios**

| **Variable** | **Coefficient** | **Hazard Ratio** | **95% CI** | **P-value** | **Significance** |
| --- | --- | --- | --- | --- | --- |
| Avg_Plas_SAtoV | 0.235 | 1.265 | 0.871 - 1.836 | 0.217 |  |
| HadPrimaryBreak | -0.747 | 0.474 | 0.286 - 0.784 | 0.004 | ** |
| T0_TLE | 0.116 | 1.123 | 1.040 - 1.212 | 0.003 | ** |
| Marker 32 | 1.911 | 6.758 | 2.777 - 16.448 | <0.001 | *** |
| W. Sambo | 0.021 | 1.021 | 0.368 - 2.834 | 0.968 |  |
| Big Pine | 0.310 | 1.364 | 0.388 - 4.795 | 0.629 |  |
| Dave's Ledge | 0.180 | 1.197 | 0.425 - 3.368 | 0.733 |  |
| Looe Key | -0.071 | 0.931 | 0.234 - 3.700 | 0.919 |  |
| EDR | 1.420 | 4.138 | 1.602 - 10.693 | 0.003 | ** |
| Maryland Shoals | 0.642 | 1.900 | 0.628 - 5.748 | 0.255 |  |
| Bahia Honda | 0.604 | 1.830 | 0.632 - 5.302 | 0.265 |  |

#### 5.3 Full Model Results

The final model incorporating interactions (n = 177 corals, 76 events) showed improved fit (concordance = 0.813, SE = 0.026) compared to the additive model and strong statistical significance (LR test: χ²(20) = 72.68, p < 0.001).

#### Table S11: Cox Model for Time to Sexual Maturity - Including Interactions.

This table presents results from the full optimized model incorporating interaction terms to examine factors affecting time to sexual maturity in *Acropora cervicornis*. The model can be expressed as:

Model: h(t) = h₀(t) × exp(β₁Site + β₂T0_TLE + β₃HadPrimaryBreak + β₄Avg_Plas_SAtoV + β₅(Site×T0_TLE) + β₆(T0_TLE×Avg_Plas_SAtoV))

Model Overview: Sample size: 177 corals Number of events: 76 corals reaching maturity Concordance: 0.813 (SE = 0.026) Likelihood ratio test: χ²(20) = 72.68, p < 0.001

| **Variable** | **Coefficient** | **Hazard Ratio** | **95% CI** | **P-value** | **Significance** |
| --- | --- | --- | --- | --- | --- |
| T0_TLE | 0.329 | 1.390 | 1.134 - 1.703 | 0.002 | ** |
| HadPrimaryBreak | -0.563 | 0.570 | 0.337 - 0.962 | 0.035 | * |
| Avg_Plas_SAtoV | 1.880 | 6.556 | 1.979 - 21.714 | 0.002 | ** |
| Marker 32 | 3.824 | 45.792 | 4.383 - 478.433 | 0.001 | ** |
| W. Sambo | -0.302 | 0.740 | 0.033 - 16.338 | 0.848 |  |
| Big Pine | 2.429 | 11.348 | 0.202 - 637.554 | 0.237 |  |
| Dave's Ledge | 0.385 | 1.470 | 0.072 - 29.816 | 0.802 |  |
| Looe Key | -2.534 | 0.079 | 0.001 - 8.828 | 0.292 |  |
| EDR | -0.047 | 0.954 | 0.073 - 12.545 | 0.971 |  |
| Maryland Shoals | 2.814 | 16.670 | 0.462 - 601.880 | 0.124 |  |
| Bahia Honda | -0.016 | 0.984 | 0.040 - 23.972 | 0.992 |  |
| T0_TLE:Avg_Plas_SAtoV | -0.177 | 0.837 | 0.738 - 0.951 | 0.006 | ** |
| Marker 32:T0_TLE | -0.219 | 0.803 | 0.625 - 1.032 | 0.086 | . |
| W. Sambo:T0_TLE | 0.041 | 1.042 | 0.764 - 1.422 | 0.796 |  |
| Big Pine:T0_TLE | -0.308 | 0.735 | 0.419 - 1.289 | 0.283 |  |
| Dave's Ledge:T0_TLE | -0.041 | 0.960 | 0.703 - 1.311 | 0.796 |  |
| Looe Key:T0_TLE | 0.252 | 1.286 | 0.855 - 1.935 | 0.226 |  |
| EDR:T0_TLE | 0.146 | 1.158 | 0.917 - 1.461 | 0.218 |  |
| Maryland Shoals:T0_TLE | -0.236 | 0.790 | 0.539 - 1.159 | 0.228 |  |
| Bahia Honda:T0_TLE | 0.050 | 1.051 | 0.765 - 1.444 | 0.758 |  |

#### 5.4 Plasticity and Post-Breakage Recovery

To test whether morphological plasticity specifically enhances post-breakage recovery rather than preventing breakage, we compared breakage rates between plasticity groups and analyzed time to sexual maturity among corals that experienced primary breaks.

#### Table S12. Breakage rates by morphological plasticity group.

Corals were classified as high (n = 105) or low (n = 153) plasticity based on median surface area to volume ratio plasticity (median = 0.87). Chi-square test showed no significant difference in breakage rates between groups (χ² = 0.12, df = 1, p = 0.734).

| **Plasticity Group** | **Total Corals** | **Corals with Any Break** | **% with Any Break** | **Corals with Primary Break** | **% with Primary Break** | **Corals with Secondary Break** | **% with Secondary Break** |
| --- | --- | --- | --- | --- | --- | --- | --- |
| High Plasticity | 105 | 86 | 81.9 | 79 | 75.2 | 7 | 6.7 |
| Low Plasticity | 153 | 129 | 84.3 | 116 | 75.8 | 13 | 8.5 |

#### Table S13. Sexual maturity outcomes for corals with primary breaks.

Among 119 corals that experienced primary branch breaks, high-plasticity individuals (n = 50) showed significantly higher maturation success (44%) compared to low-plasticity individuals (n = 69, 27.5% mature). Cox proportional hazards model: HR = 0.515 for low vs. high plasticity, p = 0.040.

| **Plasticity Group** | **N** | **Reached Maturity** | **% Mature** | **Median Time to Maturity (months)** | **Mean Time Observed (months)** |
| --- | --- | --- | --- | --- | --- |
| High Plasticity | 50 | 22 | 44.0 | 24 | 25.7 |
| Low Plasticity | 69 | 19 | 27.5 | 24 | 26.4 |

#### Table S14. Statistical test results.

Summary of statistical analyses comparing breakage rates and post-breakage maturation between plasticity groups.

| **Test** | **Test Statistic** | **p-value** |
| --- | --- | --- |
| Chi-square test: Break rates by plasticity | 0.116 | 0.734 |
| Cox model: Maturity (binary plasticity) | 24.759 | 0.003 |
| Cox model: Maturity (continuous plasticity) | 23.579 | 0.005 |
